# H3-OPT: Accurate prediction of CDR-H3 loop structures of antibodies with deep learning

**DOI:** 10.1101/2023.08.19.553933

**Authors:** Hedi Chen, Xiaoyu Fan, Shuqian Zhu, Yuchan Pei, Xiaochun Zhang, Xiaonan Zhang, Lihang Liu, Feng Qian, Boxue Tian

## Abstract

Accurate prediction of the structurally diverse complementarity determining region heavy chain 3 (CDR-H3) loop structure remains a primary and long-standing challenge for antibody modeling. Here, we present the H3-OPT toolkit for predicting the 3D structures of monoclonal antibodies and nanobodies. H3-OPT combines the strengths of AlphaFold2 with a pre-trained protein language model, and provides a 2.24 Å average RMSD_Cα_ between predicted and experimentally determined CDR-H3 loops, thus outperforming other current computational methods in our non-redundant high-quality dataset. The model was validated by experimentally solving three structures of anti-VEGF nanobodies predicted by H3-OPT. We examined the potential applications of H3-OPT through analyzing antibody surface properties and antibody-antigen interactions. This structural prediction tool can be used to optimize antibody-antigen binding, and to engineer therapeutic antibodies with biophysical properties for specialized drug administration route.

## Introduction

Antibodies protect the host by recognizing and neutralizing foreign microbes or viruses and provide immunity against future infections. Furthermore, the use of therapeutic antibodies is steadily increasing in clinical treatments for infectious diseases, cancers, and autoimmune disorders^1,2^. For example, anti-PD-L1 antibodies block PD-L1/PD-1 interactions to inhibit tumor growth in patients^3^. A typical monoclonal antibody (mAb) is composed of heavy and light chains, while a nanobody (Nb) has only a single-domain variable heavy chain. The CDR heavy chain 3 (CDR-H3) loop is the most variable region in both length and amino acid sequence, and it plays a central role in antigen binding for both mAbs and Nbs. The development of effective therapeutic antibodies requires the solved structures of candidate antibodies, including the CDR-H3. However, it is both costly and labor intensive to experimentally obtain structures, and thus computationally predicted structures are often used to guide antibody design^4^. Despite major advances in computational methods, structure prediction for CDR-H3 loops remains challenging^5^.

Traditional template-based modeling methods, such as RosettaAntibody^6^, PIGS^7^, ABodyBuilder^8^, and MODELLER^9^, often provide demonstrably inaccurate predictions for CDR-H3 sequences^10–12^, and as a result, alternative artificial intelligence (AI)-based approaches, such as AlphaFold2 (AF2)^13^, trRosetta^14^ and RoseTTAFold^15^, are increasing in popularity. AF2 has shown comparable accuracy to experimentally determined structures by capturing physical and biological information about protein folding, thus providing a versatile deep learning framework for structure prediction. Structure prediction methods using pre-trained protein language models (PLMs), e.g., HelixFold-Single^16^, OmegaFold^17^, and ESMFold^18^, have shown comparable performance to AF2 with accelerated prediction speed. PLMs can be trained with datasets comprising tens of millions of unlabeled protein sequences in a self-supervised manner, and can be subsequently applied to a variety of downstream tasks, such as druggable protein target prediction^19^, predicting protein function, and protein design^20–22^. Antibody-specific tools such as IgFold^23^, tFold-Ab^24^, DeepAb^25^ and NanoNet^26^ have also been developed to improve accuracy in CDR-H3 prediction. Among them, IgFold leverages sequence representations from PLMs to efficiently predict antibody structures within seconds, and notably, IgFold can provide accuracy comparable to AF2, enabling high-throughput prediction of antibody structures.

In this study, we present H3-OPT, which combines features of AF2 and PLMs to predict antibody structures. We compare H3-OPT with several other antibody structure prediction methods and found that it can provide a lower average RMSD_Cα_ for CDR-H3 loops than other algorithms in three subsets of varying difficulty. To further validate our model, we experimentally solved the structures of three anti-VEGF nanobodies predicted by H3-OPT^27^. We examined the potential applications of H3-OPT through analyzing antibody surface properties and antibody-antigen interactions. We demonstrate the informative value of high-quality H3 loops for predicting binding affinity, and further support the use of H3-OPT as a powerful and versatile tool for studying antigen-antibody interactions. This structural prediction tool can be used to optimize antibody-antigen binding, and to engineer therapeutic antibodies with biophysical properties.

## Results

### The H3-OPT workflow

Prior to training H3-OPT, we first conducted a thorough evaluation of currently available antibody structure prediction tools, including AF2, RoseTTAFold, trRosetta, NanoNet, DeepAb, MODELLER, and AbodyBuilder (Supplementary Section 1). AF2 provided high-accuracy predictions for the overall structures of both mAbs and Nbs, with template modeling scores (TM-scores) and global distance test scores (GDT scores) greater than 0.9. Based on the relatively higher accuracy of AF2 in structural predictions, its prediction was used to extract structural features of CDR-H3 loops for further optimization by H3-OPT. Using a mAb/Nb sequence and its Fv model structure from AF2 as input, H3-OPT was then used to generate a refined structure. As data quality has large effects on prediction accuracy, we constructed a non-redundant dataset (sequence identity < 0.8) with 1286 high-resolution (<2.5 Å) antibody structures from SAbDab^28^ (Fig. 1a). The dataset was then divided into training, validation, and test sets based on amino acid sequence identity, which was done by using the UCLUST^29^ software. To assess the prediction results, the test set was split into 3 subgroups according to the differences in AF2 accuracy (RMSD, a measure of the difference between the predicted structure and an experimental or reference structure) stemming from the length of CDR-H3 sequence: easy-to-predict targets (0 - 2 Å, Sub1); moderate difficulty targets (2 - 4 Å, Sub2); and challenging-to-predict targets (> 4 Å, Sub3), with average CDR-H3 loop sequence lengths of 9.12, 11.08, and 16.43, respectively.

**Fig. 1.**
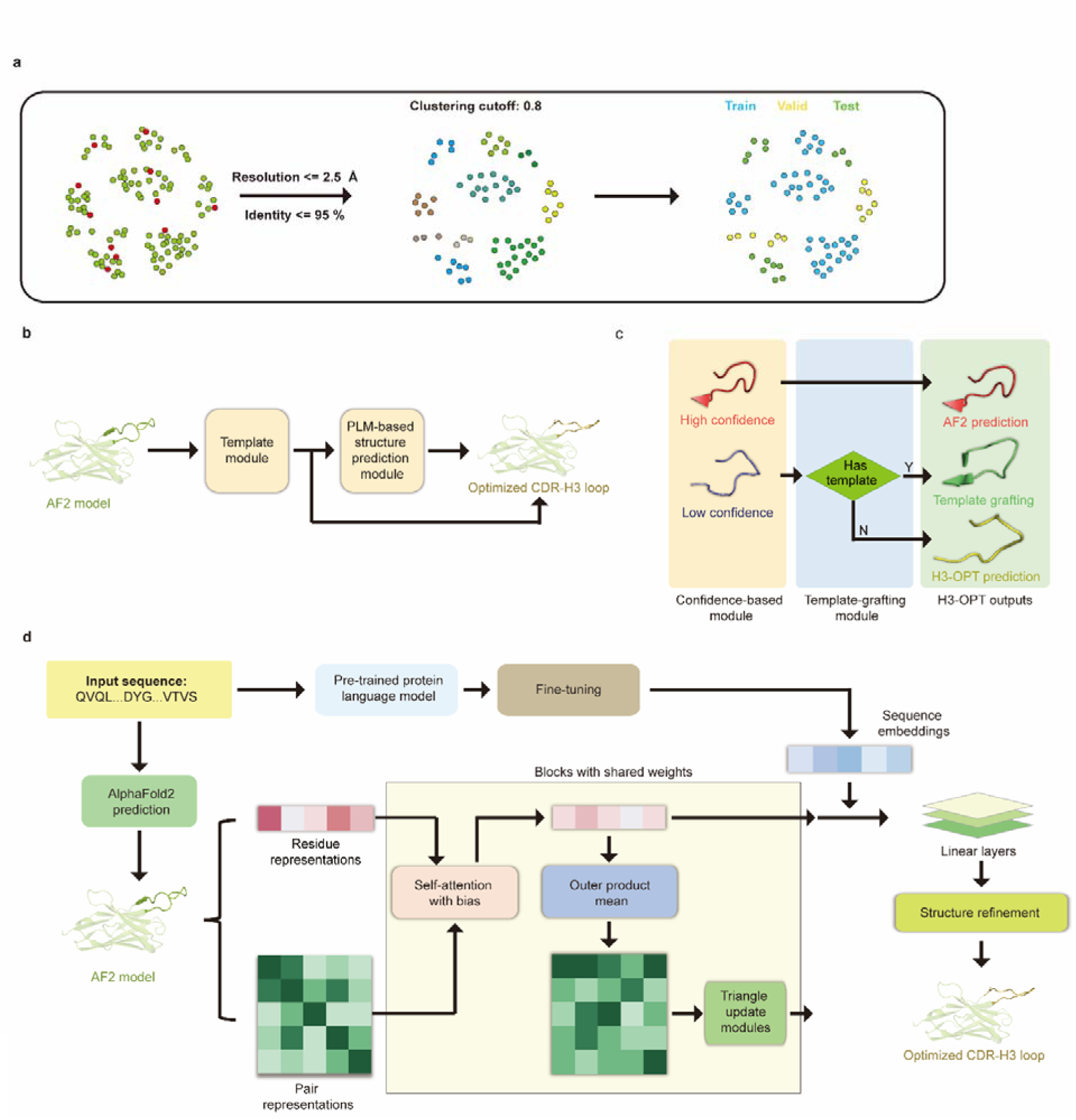
H3-OPT architecture. **a** Schematic for dataset preparation. Structures were screened from the SAbDab database based on resolution and sequence identity. Clustering of the filtered, high-resolution structures yielded three datasets for training (n = 1021), validation (n = 134), and testing (n = 131). **b,** The workflow of H3-OPT includes two modules. The template module determines whether to use PSPM, while the PSPM module optimizes the AF2 input structures. **c,** The Template module retains AF2-predicted loops when the confidence score is greater than 0.8 and grafts CDR-H3 loops onto AF2 models for structures with an available template. **d,** In the PSPM, the network extracts residue-level information and pairwise residue representations from the AF2-predicted models, which are subsequently updated using weight-sharing blocks and concatenated with sequence representations from ESM2. The resulting data is used to predict the 3D coordinates of the H3 loops.

The workflow of H3-OPT is depicted in Fig. 1b. H3-OPT consists of a Template module and a PLM-based structure prediction module (PSPM). The Template module determines whether to use PSPM to optimize CDR-H3 and comprises two sub-modules: a confidence-based module (CBM) and a template-grafting module (TGM). The CBM calculates an AF2 confidence score to evaluate the reliability of CDR-H3 loop sequence inputs through MSA and template searching. The TGM then identifies a template from the H3 template database and grafts the CDR-H3 loop onto the AF2 model if such a template is available (Fig. 1c).

The PSPM contains a series of attention-based components to ensure that the complete structural context is extracted from the input sequences. Specifically, the PSPM employs a row-wise gated attention layer that updates residue-level information and exchanges information within residue pair representations and residue features to infer relationships between the spatial and residue representations. Additionally, by leveraging sequence-level representations from PLMs along with residue-level information, the PSPM can integrate high-information features, such as protein folding patterns, across a vast protein space to predict final three-dimensional (3D) atomic coordinates (Fig. 1d).

To better understand the contribution of each module to H3-OPT accuracy with varying factors, we next examined how incorporating different approaches through the Template and PSPM modules could improve accuracy of H3-OPT. While AF2 showed markedly stronger predictions than IgFold in Sub1 (Fig. 2a), IgFold provided higher accuracy in Sub3 (Fig. 3a). In light of previous studies that showed PLM-based models are computationally more efficient but have unsatisfactory accuracy when high-resolution templates and MSA are available^18^, we therefore sought to combine the strengths of AF2 with PLM-based modeling. Similar to IgFold^23^, an initial version of H3-OPT lacking the CBM could not replicate the accuracy of AF2 in Sub1. Interestingly, we observed that AF2 confidence score of CDR-H3 shared a strong negative correlation with Cα-RMSDs (Pearson correlation coefficient =-0.67 (Fig. 2b), which led to us to hypothesize that AF2 models with high confidence scores might be sufficiently accurate, and therefore do not require further optimization. Following this notion, we incorporated a CBM sub-module that could directly retain high-confidence structures from AF2. TGM sub-module was added as a means of grafting identical H3 loop templates from public PDBs onto AF2-predicted models to further improve accuracy. Ablation studies in which the CBM or TGM were excluded to determine their respective contributions to the final predictive accuracy (Fig. 2c-f) revealed that H3-OPT without the Template Module could generate structures with average Cα-RMSDs of 1.76 Å, 2.76 Å, and 4.86 Å for the Sub1, Sub2, and Sub3 datasets, respectively.

**Fig. 2.**
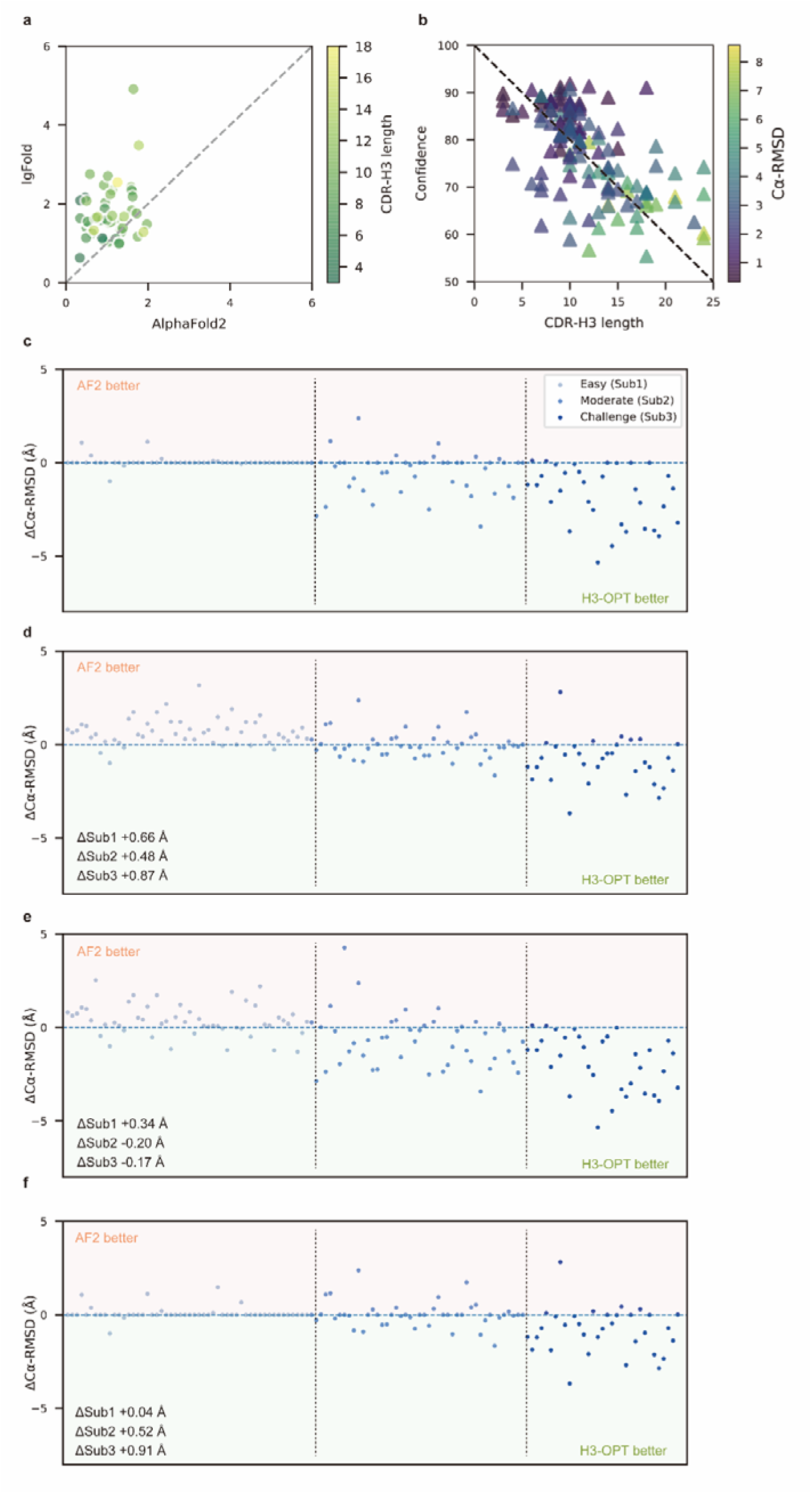
Template module and ablation studies. **a** Side-by-side comparison of Cα-RMSDs of AF2 and IgFold for Sub1 (n = 52); color scale for data points reflects CDR3 length. AF2 outperformed IgFold for targets left of the dashed diagonal; IgFold outperformed AF2 for targets right of the dashed diagonal. **b,** Correlations between AF2 confidence score and amino acid sequence length of CDR-H3 loops. Datapoint color indicates Cα-RMSD value for that target. The correlation coefficient for confidence score and CDR-H3 loop length is −0.5921. **c,** The accuracy of H3-OPT in 3 subgroups of the test set. ΔCα-RMSDs were calculated by subtracting the RMSD_Cα_ of AF2 from that of H3-OPT. AF2 had higher accuracy for targets above the dashed line; H3-OPT had better accuracy for structures below the dashed line. **d,** Differences in H3-OPT accuracy without the template module. This ablation study means only PSPM is used. **e,** Differences in H3-OPT accuracy without the CBM. This ablation study means input loop is optimized by TGM and PSPM. There are thirty targets in our database with identical CDR-H3 templates. **f,** Differences in H3-OPT accuracy without the TGM. This ablation study means input loop is optimized by CBM and PSPM.

**Fig. 3.**
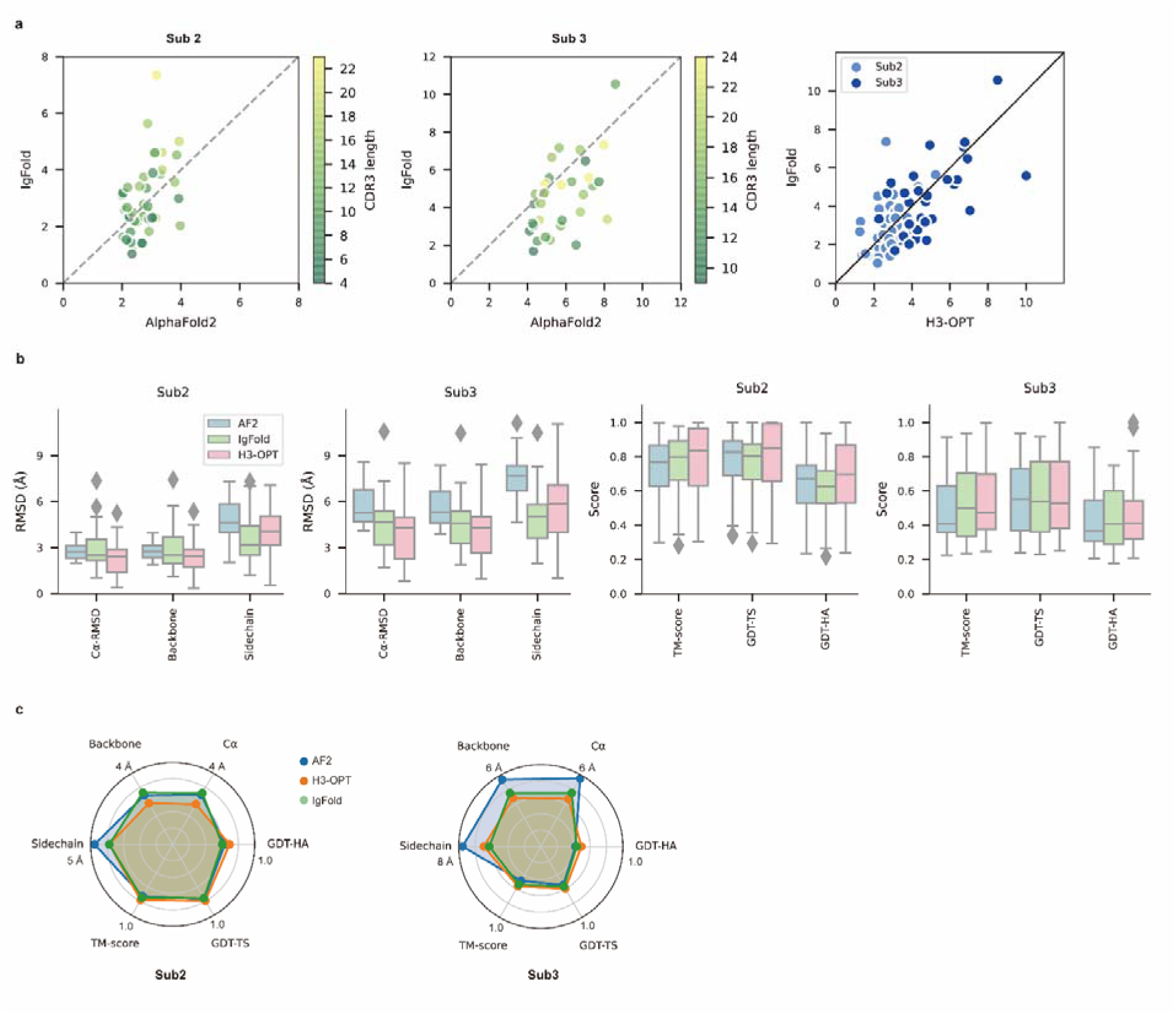
PSPM module. **a** Side-by-side comparison of Cα-RMSDs for AF2 and IgFold, IgFold and H3-OPT in the Sub2 (n = 46) and Sub3 (n = 33) test sets, respectively. **b**, Comparison of prediction accuracy between AF2 and H3-OPT for Sub2 and Sub3 targets. Metrics including RMSDs, TM-scores, and GDT-scores were used to quantitatively assess similarity between predicted and experimental structures. **c,** Comparison of prediction accuracy between AF2 and H3-OPT using six metrics (RMSD_Cα_, RMSD_backbone_, RMSD_sidechain_, TM-score, GDT-TS score and GDT-HA score). Radar plots of the mean values of different methods and metrics in predictions of Sub2 and Sub3 targets.

Incorporating only the CBM into our model significantly improved RMSD_Cα_ score by 0.62 Å for Sub 1, but exerted negligible effects on Sub2 and Sub3. By contrast, including the TGM alone resulted in substantially improved RMSD_Cα_ for Sub2 and Sub3 by 0.68 Å and 1.04 Å, respectively, suggesting that templates were more effective for relatively long CDR-H3 antibodies. These results suggested that the combination of TGM and CBM modules could leverage available templates to improve prediction accuracy.

Since the large majority of sequences in Sub2 and Sub3 have long CDR-H3 loops with few sequence homologs, attaining high accuracy in structural predictions becomes increasingly challenging for AF2^30^. Inspired by IgFold and other PLM-based methods (Fig 3a), we thus developed a PSPM module to capture structural information from the sequence embeddings of PLMs^31^. The key innovations of the PSPM for our workflow were the integration of sequence-level representations from PLMs and the simplified architecture of AF2. ProtTrans-T5, AntiBERTy, and ESM2 were initially used without fine-tuning, resulting in overall average Cα-RMSDs of 2.40 Å, 2.49 Å, and 2.32 Å, respectively (Table 1). To improve accuracy, we employed a fine-tuning approach for the downstream CDR-H3 structure prediction task. As ESM2 outperformed other PLMs in our test set, we fine-tuned parameters of all ESM2 hidden layers, which resulted in an overall RMSD_Cα_ of 2.24 Å for H3-OPT. It should be noted that most computational models, such as IgFold, froze PLM weights during model training^23^. Analysis of test subsets showed that H3-OPT could provide high accuracy in sidechain predictions, high template modeling scores (TM-scores), and high global distance test scores (GDT scores) in Sub2 and Sub3 (Fig. 3b-c).

**Table 1.** Performance of H3-OPT with different PLMs.

| | RMSDCα (Å) |
| --- | --- |
| H3-OPT | 2.24 ± 1.05 |
| AF2 | 2.85 ± 0.69 |
| ESM2 | 2.31 ± 1.13 |
| Without PLM | 2.41 ± 1.26 |
| AntiBERTy | 2.49 ± 1.42 |
| ProtTrans-T5 | 2.40 ± 1.28 |

To expand its generalizability, we simplified the Evoformer architecture and introduced parameter-sharing to directly predict Cartesian coordinates of CDR-H3 loops through both residue-level features and pair representations. In residue-level representations, rows represented amino acid types and structural features derived from AF2, while columns represented individual residues from the input sequence. The pair representations contained information about the residue pairs, such as Cα-Cα distance. These residue-level representations were updated by the attention-based layers and continuously communicated with pair representations; the pair representations were then updated via triangular multiplicative layers^30^. This simplified architecture improved model efficiency and avoided overfitting. We also applied a structural alignment strategy for feature extraction and model prediction. Given the high conservation of Fv structures, the alignment strategy was designed to effectively capture residue contribution to loop folding, resulting in improved training speed and accuracy. Taken together, H3-OPT provided high-accuracy predictions for CDR-H3 loops, facilitating the development of antibody therapeutics.

### H3-OPT provides higher accuracy CDR-H3 models than existing structure prediction software

We then evaluated the performance of H3-OPT against currently available methods, including AF2, IgFold, HelixFold-Single, ESMFold, and OmegaFold. H3-OPT achieved an average Cα root mean square deviation (RMSD_Cα_) of 2.24 Å for CDR-H3, while AF2 and IgFold had CDR-H3 RMSDs of 2.85 Å and 2.87 Å, respectively (Fig. 4a). HelixFold-Single, OmegaFold, and ESM-Fold generated comparatively poor predictions, with RMSDs for CDR-H3 of 3.39 Å, 3.75 Å, and 4.23 Å, respectively. As shown in Fig. 4b, H3-OPT provided higher accuracy predictions than other methods in all 3 subgroups, with average RMSDs of 1.10 Å, 2.28 Å and 3.99 Å, respectively. While the predictions of H3-OPT were comparable to AF2 in Sub1, its accuracy was higher than AF2 in Sub2 and Sub3.

**Fig. 4.**
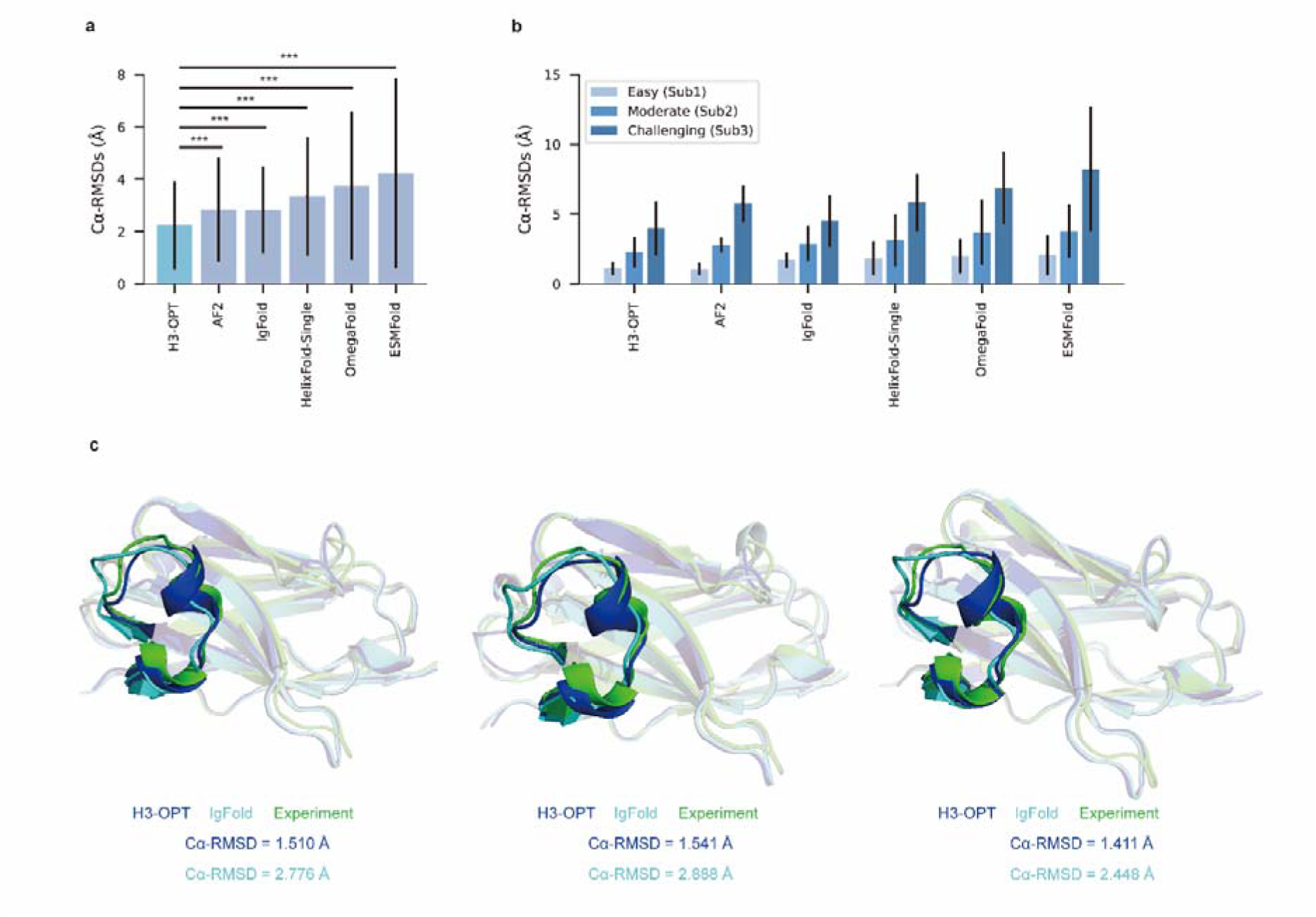
Accuracy of CDR-H3 loop prediction by H3-OPT. **a** The performance of H3-OPT in the test set (n_mAbs_ = 119, n_Nbs_ = 12) relative to other methods. The RMSD_Cα_ of H3-OPT was significantly lower than other existing methods (*p* < 0.001). **b**, The performance of H3-OPT in structural predictions of 3 subgroups of the test set (n = 52, 46, and 33). **c,** H3-OPT structural predictions for three anti-VEGF nanobodies (PDB ID: 8IIU, 8IJZ, 8IJS). The sequence identities of the VH domain and H3 loop are 0.816 and 0.647, respectively, comparing with the best template. ***, *p* < 0.001.

In light of these results, we then sought to validate H3-OPT using three experimentally determined structures of anti-VEGF nanobodies, including a wild-type (WT) and two mutant (Mut1 and Mut2) structures, that were recently deposited in Protein Data Bank (Fig. 4c).

Although Mut1 (E45A) and Mut2 (Q14N) shared the same CDR-H3 sequences as WT (Length_CDR-H3_ = 17), only minor variations were observed in the CDR-H3. H3-OPT generated accurate predictions with Cα-RMSDs of 1.510 Å, 1.541 Å and 1.411 Å for the WT, Mut1, and Mut2, respectively (The confidence scores of these AlphaFold2 predicted loops were all higher than 0.8, and these loops were accepted as the outputs of H3-OPT by CBM). Subsequent comparison with IgFold, showed that AlphaFold2 outperformed IgFold on these targets and IgFold could not accurately predict the short helix in CDR-H3, resulting in Cα-RMSDs of 2.776 Å (WT), 2.888 Å (Mut1), and 2.448 Å (Mut2) and more diverse conformations of Mut1 and Mut2. These results indicated that IgFold was capable of learning long-range correlations in protein sequences, but that these long-range correlations could introduce larger errors when MSA and templates were available. These results thus demonstrated that MSA could strongly influence the accuracy for which similar CDR-H3 template structures were available.

### H3-OPT can predict antibody surface properties

To demonstrate how improved CDR-H3 structural accuracy can assist antibody engineering, we applied H3-OPT to predict surface properties. Comparison of H3-OPT with AF2 for identifying surface amino acids (SAAs) by the relative accessible surface areas (rASAs) of CDR-H3 loops showed that H3-OPT could predict SAAs, on average, close to that of the native structures (9.46 vs 9.40, Fig. 5a), whereas AF2 predicted an average of 9.81 SAAs. Furthermore, H3-OPT predicted closer values to native structures than AF2 in predicting diverse surface properties, such as the distribution of hydrophilic or charged SAAs (Fig. 5b).

**Fig. 5.**
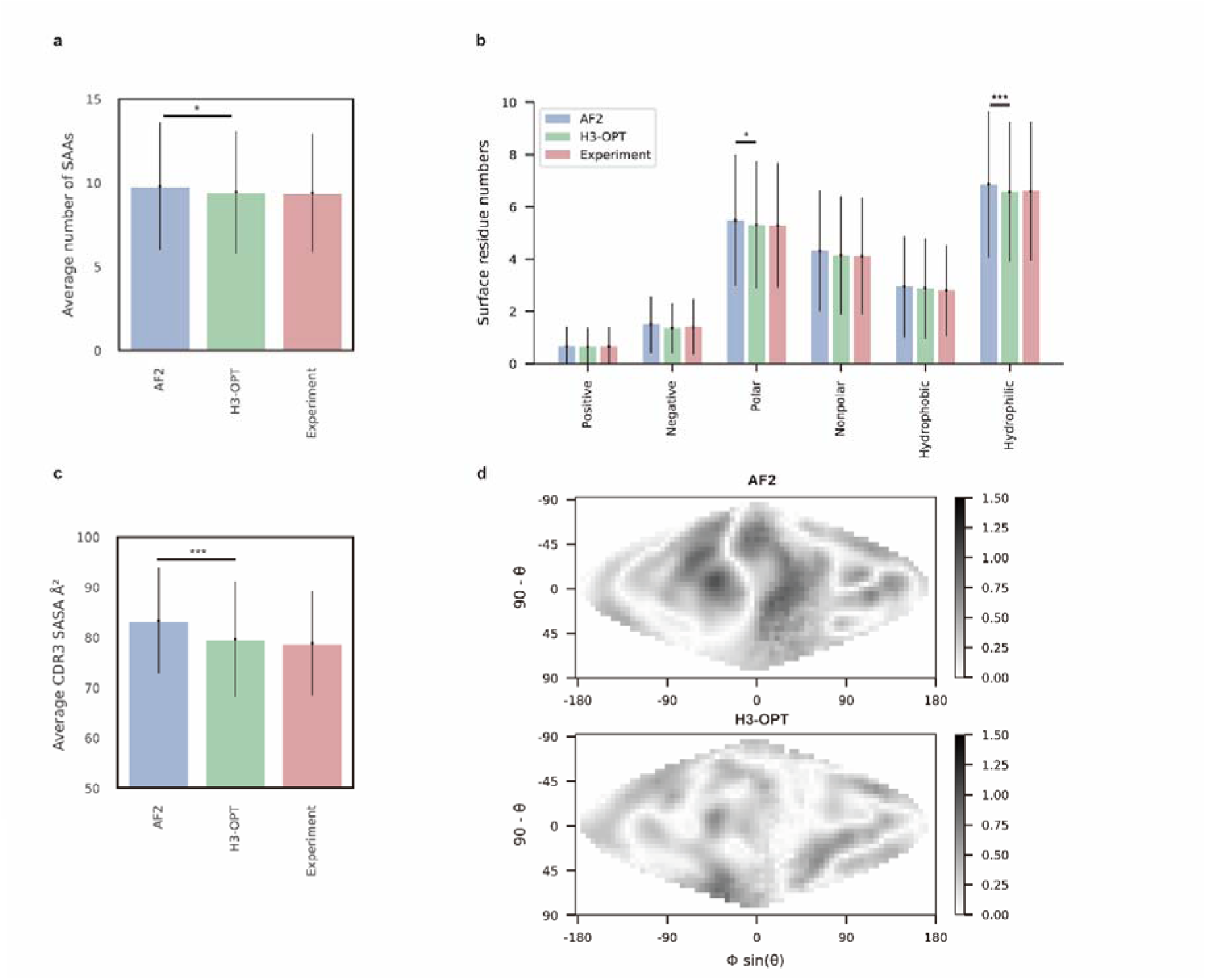
Analysis of surface patches. **a** Analysis of surface amino acids for predicted H3 loops. Y-axis represents average number of surface residues for H3 loops (n = 131). The surface residues of AF2 models are significantly higher than those of H3-OPT models (p < 0.05). **b,** Histogram of surface patches with different properties predicted by H3-OPT, AF2, or experimentally solved H3 loops. Error bars show standard deviations. H3-OPT models predicted lower values than AF2 models in terms of various surface properties, including polarity (p <0.05) and hydrophilicity (p < 0.001). **c,** Solvent-accessible surface area (SASA) analysis of predicted H3 loops. Values represent the difference in SASA between predicted and experimentally determined H3 structures using AF2 or H3-OPT. The SASA of AF2 models are significantly higher than those of H3-OPT models (p < 0.001). **d,** Comparison of the charged surface patches between H3-OPT and AF2 for target PDBID: 5U3P. The surface maps compare the surface electrostatic potential of the CDR-H3 loop predicted by H3-OPT or AF2 with the native structure. Darker shading indicates greater difference in electrostatic potential. *, *p* < 0.05; **, *p* < 0.01; ***, *p* < 0.001.

To examine insights H3-OPT could provide into the biophysical properties of antibodies^32^, we next estimated the solvent-accessible surface areas (SASAs). The average SASAs of H3-OPT loops closely resembled that of the native structures, with larger differences in AF2 predictions (Fig. 5c). Plots of SASA distributions at each alignment position revealed smaller errors in H3-OPT than AF2, with ΔSASA ranging from -12.17 to 9.74 Å^2^ (Extended Data Fig. 1). In addition, comparison of surface charge distributions on the predicted CDR-H3 loops generated by H3-OPT and AF2 in an electrostatic map of a representative structure (PDB 5U3P, Fig. 5d) showed that H3-OPT predictions were consistent with the experimental structure. These results collectively showed that accurate surface properties predicted by H3-OPT could provide insights into antibody folding and stability, as well as their interactions with antigens.

### H3-OPT can facilitate investigation of antibody-antigen interactions

To explore the potential applications of H3-OPT in studying antibody-antigen interactions, we analyzed contact patterns at predicted binding sites. Given that predicting the structure of an antibody-antigen complex remains challenging, we superimposed V_H_ fragments from H3-OPT and AF2 onto experimentally determined native complex structures available in our test set. After identifying the contact residues of antigens by H3-OPT, we found that H3-OPT could substantially outperform AF2 (Fig. 6a), with a median precision of 0.82 and accuracy of 0.98 compared to 0.71 precision and 0.97 accuracy of AF2. Next, we estimated the distances among interface residues, which are related to binding affinity at the interface. We found that H3-OPT had less error in distance metrics than predictions by AF2 across different distance thresholds, with average mean squared errors of 2.42 Å and 4.85 Å, respectively (Fig. 6b). Furthermore, calculation of H3 contact propensities to assess potential differences in binding patterns between the predicted and native structures using high-quality H3 conformations indicated that H3-OPT also displayed higher accuracy in predicting contact propensities (Fig. 6c).Finally, to test whether H3-OPT was reliable for investigation of the detailed mechanisms underlying antigen-antibody binding, we generated contact maps and calculated binding affinities for complex structures obtained by H3-OPT, AF2, or through experiments (see Methods for details). We found that contact maps predicted by H3-OPT were consistent with those observed in the experimental structures (Fig. 6d). Since affinity prediction plays a crucial role in antibody therapeutics engineering, we performed MD simulations to compare the differences in binding affinities between AF2-predicted complexes and H3-OPT-predicted complexes. Calculation of binding affinities through MD simulations showed that the average affinities of structures obtained by H3-OPT prediction were closer to those of experimentally determined structures than values obtained through AF2 (Table 2). These cumulative findings illustrate the informative value of high-quality H3 loops for predicting binding affinity and facilitating antibody engineering, and further support the use of H3-OPT as a powerful and versatile tool for investigating antigen-antibody interactions.

**Fig. 6.**
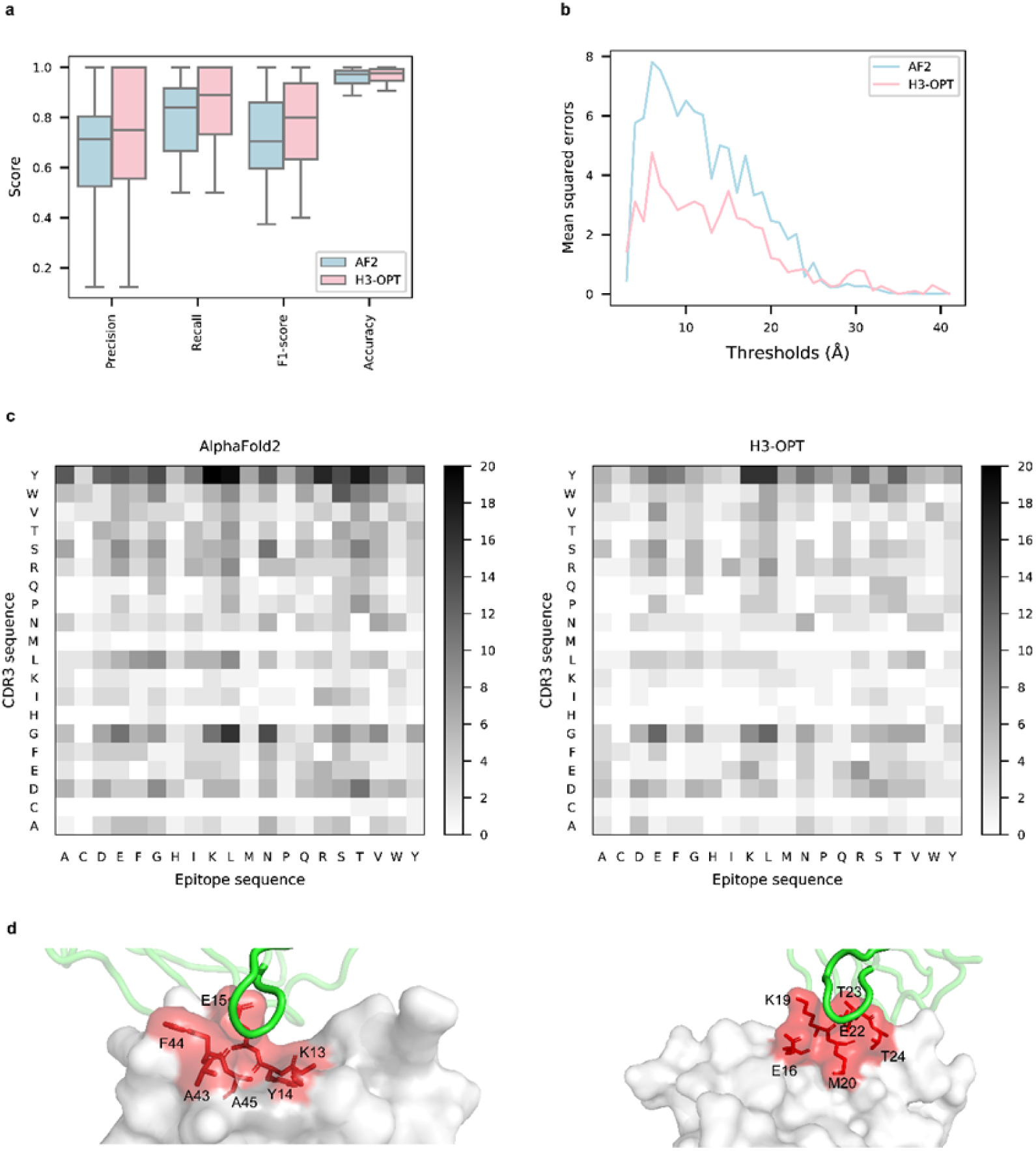
Accuracy of H3-OPT predictions of antibody-antigen interactions. **a** Performance of H3-OPT in binding site prediction. comparison of prediction accuracy between H3-OPT and AF2 for antibody-antigen binding sites (n = 27). Box represents interquartile range (IQR); horizontal line in the center of the box shows median. **b,** Comparison of the mean squared errors of residue pairs between H3-OPT and AF2 under different distance thresholds. The x-axis represents the experimentally determined distance between pairs of contacting residues at the binding site in the native structure. Y-axis shows mean squared errors of H3-OPT and AF2. **c,** Heatmaps of the frequency of pairwise residue-residue contacts across antibody-antigen interfaces. This analysis compares contact frequency of H3 loops predicted by AF2 or H3-OPT with the native structure. Darker shading indicates greater difference in contact frequency. **d,** The predicted H3 loops of two targets interacting with antigens (PDB: 2YC1, 6O9H). The epitopes are highlighted in red and antibody chains are green. H3-OPT could identify the epitopes of different antigens that form the complementary binding interface(s) for the CDR-H3 of antibodies.

**Table 2.** Comparison of binding affinities obtained from MD simulations using AF2 and H3-OPT.

| PDBID | AF2 (kcal/mol) |  |  | H3-OPT (kcal/mol) |  |  | AF2 (kcal/mol) |  | H3-OPT (kcal/mol) |  |
| --- | --- | --- | --- | --- | --- | --- | --- | --- | --- | --- |
| | MM/GBSA | MM/PBSA | AF2 RMSD<sub>Ca</sub> (Å) | MM/GBSA | MM/PBSA | H3-OPT RMSD<sub>Cα</sub> (Å) | $\vert \Delta_{MM/GBSA}\vert $ | $\vert \Delta_{MM/PBSA}\vert $ | $\vert \Delta_{MM/GBSA}\vert $ | $\vert \Delta_{MM/PBSA}\vert $ |
| 2ghw | -29.20 | -33.36 | 2.7 | -14.70 | -21.36 | 3.0 | 8.63 | 2.42 | 23.13 | 14.42 |
| 2yc1 | -38.85 | -37.73 | 2.3 | -43.80 | -48.72 | 1.5 | 6.80 | 18.67 | 1.85 | 7.68 |
| 3l95 | -29.59 | -53.35 | 2.5 | -47.22 | -68.86 | 2.5 | 23.60 | 11.44 | 5.97 | 4.07 |
| 3u30 | -37.31 | -42.41 | 2.6 | -44.94 | -50.07 | 2.5 | 9.64 | 2.18 | 2.01 | 5.48 |
| 4cni | -36.96 | -42.93 | 1.0 | -31.92 | -40.39 | 1.3 | 8.54 | 7.44 | 3.50 | 4.89 |
| 4nbz | -36.59 | -43.79 | 1.9 | -59.61 | -54.23 | 0.6 | 10.17 | 3.30 | 12.85 | 13.74 |
| 4xnq | -13.55 | -17.47 | 2.7 | -31.51 | -30.94 | 0.5 | 15.40 | 12.32 | 2.57 | 1.15 |
| 4ydl | -52.51 | -74.25 | 4.8 | -49.17 | -73.57 | 3.6 | 6.82 | 6.53 | 10.15 | 7.21 |
| 5e5m | -59.72 | -71.15 | 3.0 | -41.29 | -53.70 | 7.3 | 0.50 | 5.79 | 18.93 | 11.66 |
| 5f7y | -61.76 | -69.46 | 2.7 | -60.33 | -69.43 | 1.4 | 3.95 | 6.38 | 5.38 | 6.41 |
| 6kyz | -12.66 | -20.32 | 4.0 | -9.36 | -17.13 | 3.7 | 17.63 | 17.21 | 20.93 | 20.40 |
| 6o9h | -39.53 | -43.45 | 2.8 | -52.27 | -57.51 | 0.6 | 10.45 | 13.31 | 2.29 | 0.74 |
| 6pyd | -45.87 | -58.71 | 1.0 | -35.75 | -45.28 | 1.1 | 6.29 | 13.50 | 3.83 | 0.06 |
| 6u9s | -36.54 | -48.66 | 1.0 | -39.79 | -44.80 | 1.3 | 14.35 | 10.42 | 11.11 | 14.28 |
| Average | / | / | 2.6 | / | / | 2.4 | 10.20 | 9.35 | 8.89 | 8.01 |

\* $\Delta_{MM/GBSA}$ (or $\Delta_{MM/PBSA}$ ) was calculated by subtracting the MM/GBSA (or MM/PBSA) of predicted model from the MM/GBSA (or MM/PBSA) of experimental structure.

## Discussion

Several platforms for in silico antibody engineering have been developed to improve binding affinity^33–36^, humanization^37,38^ and stability^39,40^. At present, engineering these biophysical properties of therapeutic antibodies heavily relies on the availability of the structures of thousands of antibodies. The accurate structure of CDR-H3 is crucial for understanding the mechanism(s) of antigen-antibody interactions. Although AF2 performs well when MSA and templates are available, its predictive capability decreases for targets with long CDR-H3 loops. By contrast, PLMs leverage tens of millions of protein sequences to learn relevant structural patterns, consequently enhancing their capacity to predict orphan antibodies^41^. The results of IgFold indicate that PLMs, which incorporate transformer-based attention mechanisms, are capable of learning both local and global sequence information^23^. While global information can be useful for challenging targets, it may lead to large errors when templates were available, as demonstrated by our case study of anti-VEGF nanobodies (Fig. 4).

H3-OPT combines the strengths of AF2 and PLMs to predict the structures of mAbs and Nbs, and our results show that combining these features can improve the average RMSD_Cα_ by 20% over that of AF2 or IgFold. However, H3-OPT is less efficient than PLM-based methods like IgFold because it relies on AF2 structures. To improve efficiency, one possible solution is to cluster the targets based on sequence identity and compute AF2 structures for each cluster, followed by generation of mutant structures. Despite the lower efficiency, H3-OPT offers several notable advantages over IgFold. First, H3-OPT incorporates a simplified version of Evoformer, which improves training efficiency. In addition, H3-OPT employs an alignment-based strategy to extract structural features that are then used to directly predict the Cα coordinates of H3 loops, effectively capturing the residue features relevant to loop folding. Moreover, we found that IgFold uses an augmented dataset containing AF2-predicted structures, which could potentially propagate errors during model training. In contrast, H3-OPT was trained by using high-quality and non-redundant antibody structures. Finally, H3-OPT also shows lower Cα-RMSDs compared to AF2 or tFold-Ab for the majority (six of seven) of targets in an expanded benchmark dataset, including all antibody structures from CAMEO 2022 (Extended Data Fig.2).

Although H3-OPT can provide high accuracy with a small training dataset, it is likely that a larger dataset containing high-resolution structures will further improve our model. In the current study, we observed that the accuracy of AF2 CDR-H3 predictions was correlated with the length of the H3 loop. Thus, AF2 can provide accurate predictions for short loops with fewer than 10 amino acids, with PLM-based models offering little or no improvement in such cases. Conversely, antibodies with H3 loop lengths exceeding 25 residues (common in Sub3) pose a long-standing challenge for structural prediction algorithms, and none of the existing methods (including H3-OPT) tested here could provide accurate predictions due to a lack of homologous sequences and a high degree of freedom in this subgroup. Notably, H3-OPT outperformed AF2 in Sub3 because the context of protein sequences learned from ESM2 was fine-tuned to fit antibody structures. Furthermore, we envision that antibodies with H3 loop lengths ranging from 10 to 25 amino acids could be further optimized using PLMs with more model parameters. In addition, we attempted to optimize the H3 loop structure through molecular dynamics simulations and quantum mechanics-based methods (Supplementary Section 2). However, these approaches generally yielded less accurate predictions than AF2 due to inaccurately described solvent effects and other environmental factors. The development of a more accurate, physics-based method for modeling long loops may be another potentially effective strategy for improving H3-OPT.

Deep generative models are playing a vital role in the field of protein design, leading to several successful applications^42–47^. These models capture the intricate relationship between sequences and structures, enabling the generation of novel protein scaffolds not existing in nature. However, the potential immunogenicity concerns associated with these newly designed proteins require comprehensive preclinical and clinical investigations. Consequently, conventional mAbs and Nbs remain the prevailing choices in medicine. Recognizing that the properties of antibodies are heavily influenced by their structures, accurate prediction of the structures of mAbs and Nbs is of great significance in optimizing their therapeutic effectiveness and clinical relevance. To this end, H3-OPT has strong potential to accelerate optimization of antibody-antigen binding as well as engineering of therapeutic antibodies with specialized biophysical properties.

## Method

### H3-OPT dataset

Crystal structures of antibodies were obtained from SAbDab, a non-redundant nanobody database^48^ and a subset of DeepAb. The structures with resolution greater than 2.5 Å or redundant sequences with 95% or greater identity were removed (Fig. 1a). We removed the structures with missing loops in CDR-H3 or lengths greater than 45. The structures with missing residues at CDR-H3 loops were also dropped. To enhance the generalizability of H3-OPT, the sequences were clustered based on a 90% similarity cutoff by using UCLUST^29^. The resulting clusters were then randomly divided into training, validating, and testing sets to avoid overestimation of performance. The number of sequences in training, validating and testing sets was 1021(925 mAbs, 96 Nbs), 134 (122 mAbs, 12Nbs) and 131 (119 mAbs, 12 Nbs), respectively, with average CDR-H3 loop length of 11.8, 11.9 and 11.7, respectively. Schrödinger v. 2017-2 (Schrödinger, New York, NY, USA), Numpy 1.18.5 and Pandas 1.0.5 was used for data preparation.

### Structure preparation

To address the transformational invariance for predicting 3D coordinates, we applied a structural alignment strategy. Prior to feature extraction, we removed all atoms of light chains in mAbs Fv structures and any non-standard residues in native structures. All antibody V_H_ native structures were aligned using StructAlign module in Schrödinger with a randomly selected reference structure (PDBID: 1GIG). Next, we superimposed AF2 predicted models to their corresponding native structures. This alignment allowed the model to learn the interactions between residues located in H3 loops while disregarding the rotation and translation of the entire structure. The heavy chain fragments were determined using Chothia definitions^49^.

### Feature generation

As shown in Table 3, structural features were extracted and aggregated into the following inputs to PSPM of H3-OPT: A residue feature vector of size [N_res_, 34] was constructed by concatenating “Amino acid type”, “AF2 predicted coordinates”, “AF2 predicted backbone torsion angles”, “Torsion angles mask” and “Predicted residue mask”. The pairwise feature vector of size [N_res_, N_res_, 60] included the “Pairwise distances” and “Pairwise amino acid type”.

**Table 3.**
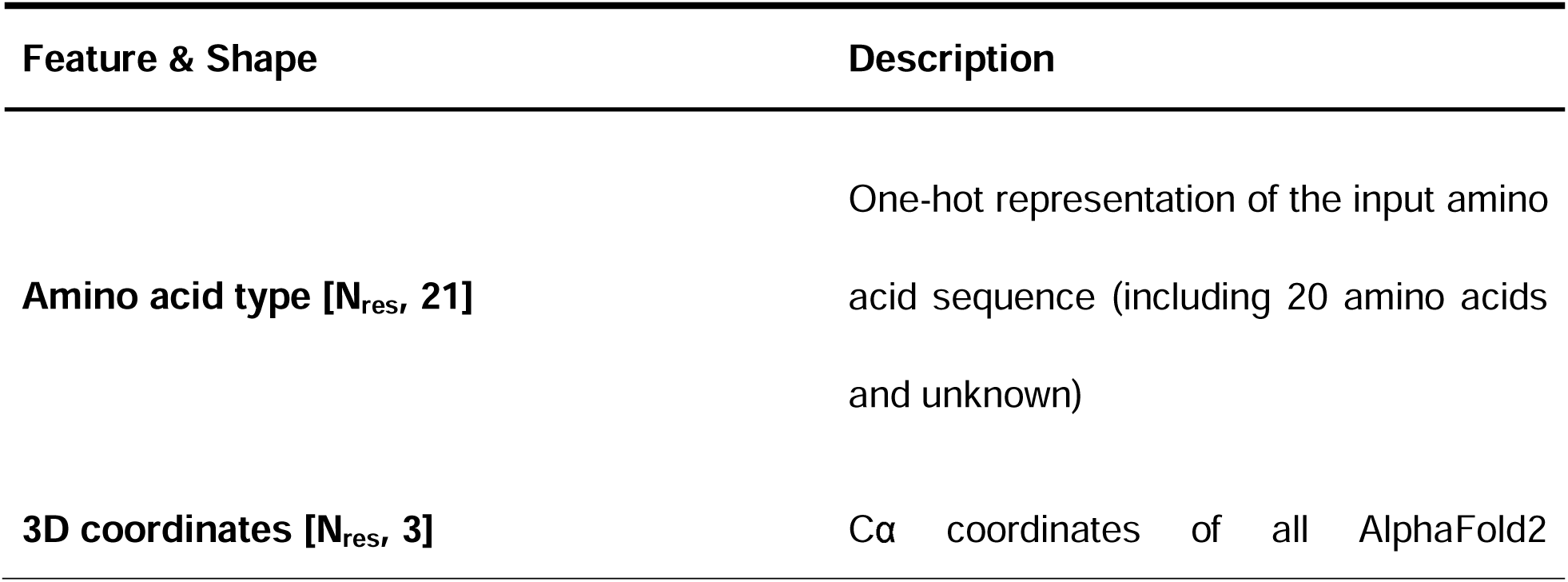
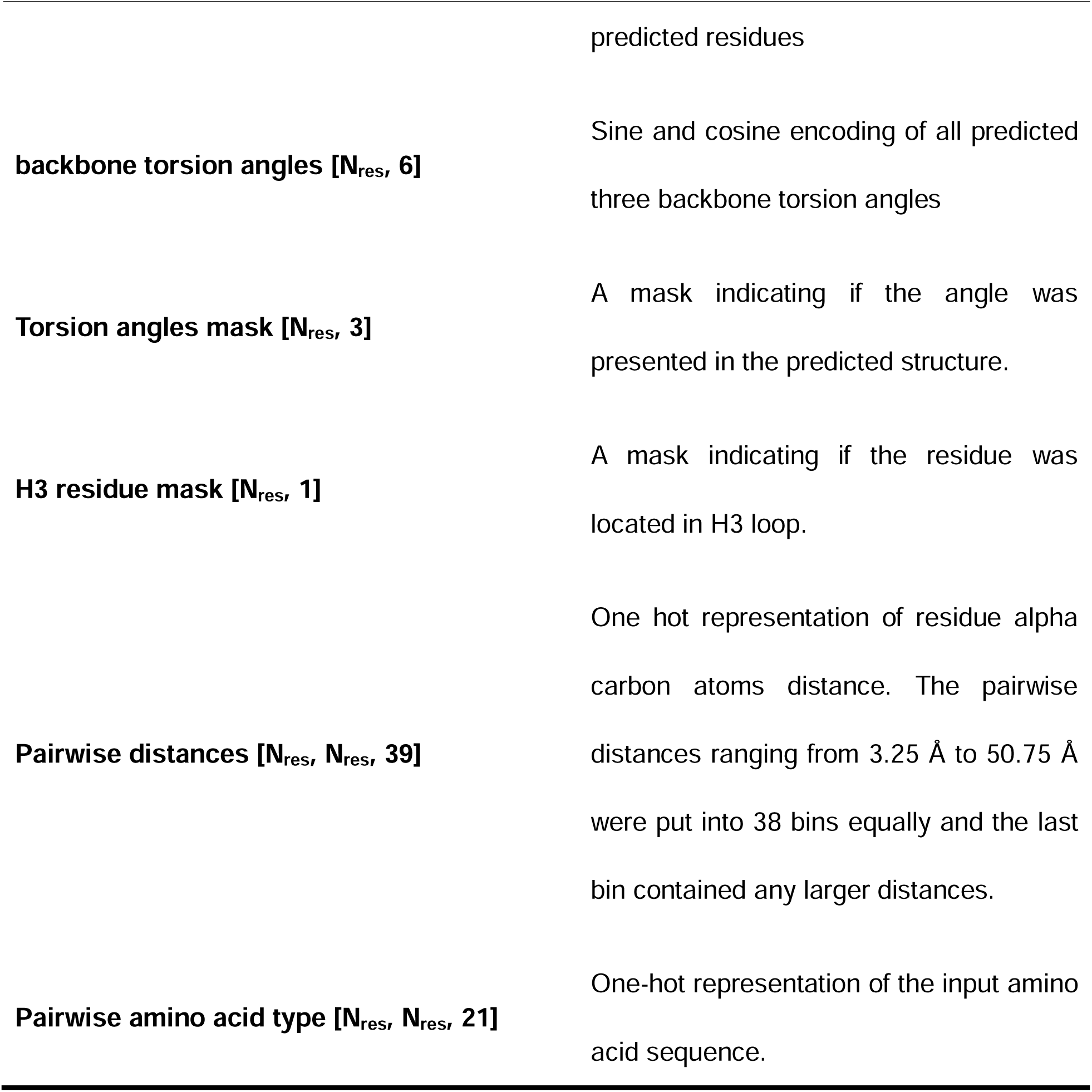
Features to the model. N_res_ is the number of residues. ^30^

### Network architecture

H3-OPT took the predicted H3 loops of AF2 as input and generate the antibody structure with the optimized CDR-H3 loops as output. H3-OPT was composed of a Template module and a PSPM. The Template module contains two submodules, i.e., CBM and TGM. The CBM calculated the average confidence score of H3 residues and retained AF2 structures with confidence score greater than 0.8. The TGM grafted the template loop structure onto the input AF2 model, when their CDR-H3 sequences were identical. We replaced the predicted CDR-H3 loop generated by AF2 with the template loop by aligning corresponding atoms, which included Cα atoms and carbonyl carbon atoms in the first two residues and Cα atoms and backbone nitrogen atoms in the last two residues.

The input residue-level features are extracted from AF2 predicted structures as mentioned above, resulting in a dimension of *L* × 34, where L is the padding length of antibody heavy chain. Similarly, the pairwise features has a dimension of *L* × *L* × 60. The PSPM network began with a row-wise gated multi-head self-attention layer which updated residue-level features from pair representations. After that, the residue-level information updated the pair representations via outer product mean module. Pair representations were updated by two multiplicative update modules. During training, the weights of the attention layer and the triangle multiplicative update modules were updated 4 times per epoch. Then, the final hidden states from ESM2 (esm2_t33_650M_UR50D) were passed to a linear layer and then concatenated with the residue-level features to enable the learning of latent information from vast protein sequences. To predict all Cα coordinates for the CDR-H3 loop, the output vector was passed through three linear layers with a hidden size of 64. In summary, PSPM utilized an attention-based mechanism to learn the contribution of individual residues and incorporate sequence representations from PLMs into final predictions. We used OpenFold^50^ (based on GitHub from June 2022: commit 3f57b4a041f063406059f42080ede6d495479617) to implement partial networks of AF2.

We employed a fine-tuning strategy to update the sequence representations in ESM2. Initially, we froze all weights of the 33 representation layers in ESM2 and updated only the weights of the attention layers and pair representation update modules. Subsequently, we fixed all weights of the remaining components in H3-OPT and performed fine-tuning exclusively on all hidden layers of ESM2 for the H3 loop prediction task. Finally, we fine-tuned all parameters of H3-OPT to predict the atomic coordinates of all CDR-H3 Cα atoms.

During training, we trained our model on the training set and validated it on the validation set. The dropout rates of H3-OPT were set to 0.25. Mean squared error (MSE) loss was utilized to train the model. We used Adam optimizer with a learning rate of 1LJ×LJ10^−4^, weight decay of 5LJ×LJ10^−4^ for training. The model was trained on an NVIDIA V100 super GPU and took one hour per epoch over the entire training set with a batch size of 64. We randomly selected 10% of all structures for model validation. The H3-OPT was implemented using PyTorch 1.12.1 in Python 3.7.2. The hyperparameters used during training process for H3-OPT were presented in Table 4.

**Table 4.**
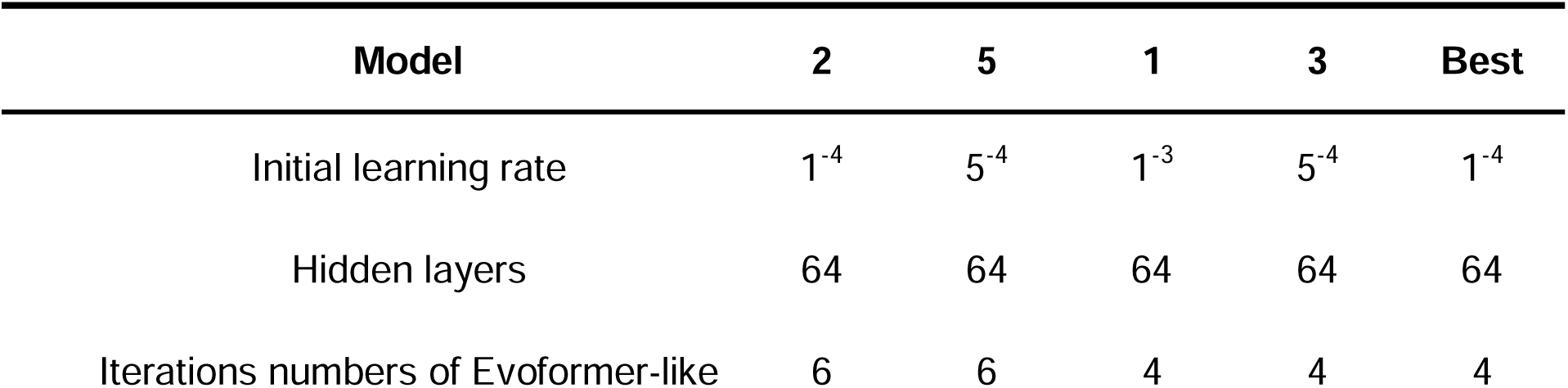
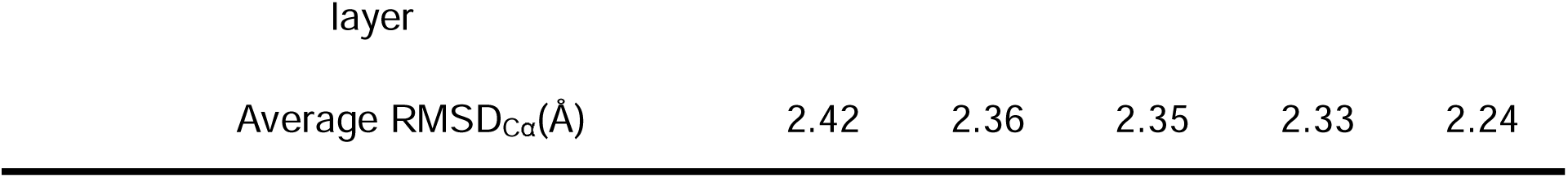
Hyperparameters for H3-OPT models.

**Table 5.** Average Cα-RMSDs of our test set under different confidence cutoffs.

| Cutoff | Cα-RMSD (Å) |
| --- | --- |
| 0.70 | 2.46 |
| 0.75 | 2.30 |
| 0.80 | 2.24 |
| 0.85 | 2.17 |
| 0.90 | 2.29 |
| 0.95 | 2.28 |

### Structure generation and refinement

We utilized a structure refinement strategy to generate structures of CDR-H3 loops and rectify structural errors. Initially, we used PSPM predictions to modify the Cα coordinates of predicted CDR-H3 loops generated by AF2. Then, we minimized the remaining atoms of the CDR-H3 loops, while keeping their Cα coordinates fixed, by using the OPLS 2005 force field^51^. After the initial energy minimization step, we applied a force constant of 10 kcal/mol to these fixed atoms to guarantee complete relaxation of the entire loops.

### Validation settings

H3-OPT, AF2, HelixFold-Single, IgFold and ESMFold were compared on the test subset. Given that tFold-Ab is only available on webserver, we compared H3-OPT with tFold-Ab on CAMEO 2022 dataset. We removed the target antibody structures from AF2 database before prediction to evaluate the performance of AF2. This ensured that the predictions depended solely on the information derived from other template structures, thus avoiding biased results. Since all other methods did not require MSA searching, we did not exclude these targets from their database. To predict the entire Fv structures, we included 50 glycine residues as a linker between the heavy chain and the light chain for antibodies^52^. The Cα-RMSDs of all predicted H3 loops were calculated after superimposing the V_H_ backbone heavy atoms (without CDR-H3) to the reference structures. All structures were generated using publicly available code repositories: AF2 v2.1.1(based on GitHub the November 2021 version at GitHub: commit 91b43223422420d1783ed802c8b3a8382a9309fd), IgFold (based on GitHub from February 2023: commit 6a09298d165ed1deb438c0b6eefcbcb03ed0eca5), HelixFold-Single (based on GitHub from January 2023: commit 5f39b2c2a4ecc00b89ba05b95dc56212bdd5d886) and ESMFold v1 (based on GitHub from October 2022: commit dc823b89c6acb9f67caea53704c2a97524fbd456).

### Analysis of SASAs and pairwise residue contacts

We computed the SASAs and rASAs of all CDR-H3 loops by using the ANARCI webserver^53^ with the default settings. Residue with an rASA greater than 25% was considered surface residues^54^. Subsequently, we applied AHo alignment software^55^ to report the averaged SASA per alignment position. The samples and alignment positions with more than 10 gaps were removed to avoid randomness and bias, reducing the final samples to 123. Finally, the difference in SASAs between H3-OPT and AF2 was calculated by subtracting predicted H3 loops SASAs from the ground truth.

To examine the contribution of H3-OPT in discovering antibody-antigen interactions, we identified contact residue pairs that were within this distance as binding sites by setting a threshold of 5 Å. We next calculated the pairwise residue distance matrices for each individual predicted complex and native structure, where each element represented the closest distance between the heavy atoms of two residues. From these pairwise distances, we derived contact propensity matrices that specifically indicated the presence or absence of interactions between residues.

### Analysis of surface electrostatic potential

We generated two-dimensional projections of CDR-H3 loop’s surface electrostatic potential using SURFMAP^56^ v2.0.0 (based on GitHub from February 2023: commit: e0d51a10debc96775468912ccd8de01e239d1900) with default parameters. The 2D surface maps were calculated by subtracting the surface projection of H3-OPT or AF2 predicted H3 loops to their native structures.

### Binding affinity calculation

We analyzed the binding affinities of antibody-antigen complexes predicted by AF2 and H3-OPT. Their relative binding affinities were calculated through molecular dynamics simulations. Initially, the missing side-chains and loops in the antibody structures were filled using the Protein Preparation Wizard module within Schrödinger software. Subsequently, the tLEaP module of AMBER^57^ was employed to construct the simulation system with the ff19SB force field^58^ and OPC solvent model^59^. Additionally, the simulation system was solvated with a 0.15 M NaCl solution. Energy minimization was performed through a 5000-step steepest descent algorithm, followed by a 5000-step conjugate gradient algorithm. Afterwards, a 400-ps NVT simulation with a time step of 2 fs was performed to gradually heat the system from 0 K to 298 K (0–100 K: 100 ps; 100-298 K: 200 ps; hold 298 K: 100 ps), and a 100-ps NPT simulation with a time step of 2 fs was performed to equilibrate the density of the system. During heating and density equilibration, we constrained the antigen-antibody structure with a restraint value of 10 kcal·mol^-1^·Å^-2^. In the production run, 100-ns MD simulations were performed with a time step of 2 fs. The first 50 ns restrains the non-hydrogen atoms of the antigen-antibody complex, and the last 50 ns restrains the non-hydrogen atoms of the antigen, with a constraint value of 10 kcal·mol^-1^·Å^-2^. The distance cutoff for nonbonded interactions was set to 10 Å, and the Berendsen algorithm was utilized to maintain isotropic pressure coupling at 1 bar. The Langevin algorithm was employed to maintain the simulation temperature at 298 K. The relative binding affinities of the antigen-antibody complexes were evaluated using the MMPBSA module of AMBER software, which computed the MM/GBSA energies for the trajectory frames of last 10 ns.

## Crystallization and data collection

The protein expression, purification and crystallization experiments were described previously^27,60^. The proteins used in the crystallization experiments were unlabeled. Upon thawing the frozen protein on ice, we performed a centrifugation step to eliminate any potential crystal nucleus and precipitants. Subsequently, we mixed the protein at a 1:1 ratio with commercial crystal condition kits using the sitting-drop vapor diffusion method facilitated by the Protein Crystallization Screening System (TTP LabTech, mosquito). After several days of optimization, single crystals were successfully cultivated at 21°C and promptly flash-frozen in liquid nitrogen. The diffraction data from various crystals were collected at the Shanghai Synchrotron Research Facility and subsequently processed using the aquarium pipeline.^61^

## Statistics analysis

We conducted two-sided t-test analyses to assess the statistical significance of differences between the various groups. Statistical significance was considered when the p-values were less than 0.05. These statistical analyses were carried out using Python 3.10 with the Scipy library (version 1.10.1).

## Data and code availability

The datasets of our study and the codes of H3-OPT are freely available at https://github.com/chdcg/H3-OPT.

## Acknowledgments

This work was supported by the Tsinghua University Initiative Scientific Research Program (No.20231080030), Vanke Special Fund for Public Health and Health Discipline Development (2022Z82WKJ009), the Tsinghua-Peking University Center for Life Sciences (No.20111770319), Tsinghua University - Peking Union Medical College and Hospital Collaboration Foundation (No. 20191080837).

## Author contributions

H.C. and B.T. conceptualized the project. H.C. and X.F. developed the methodology. S.Z. solved crystal structures. H.C. and L.L. performed structural modeling. H.C. wrote the original paper. Q.F., X.Z., and B.T. edited the paper.

---

**Extended Data Fig. 1.**
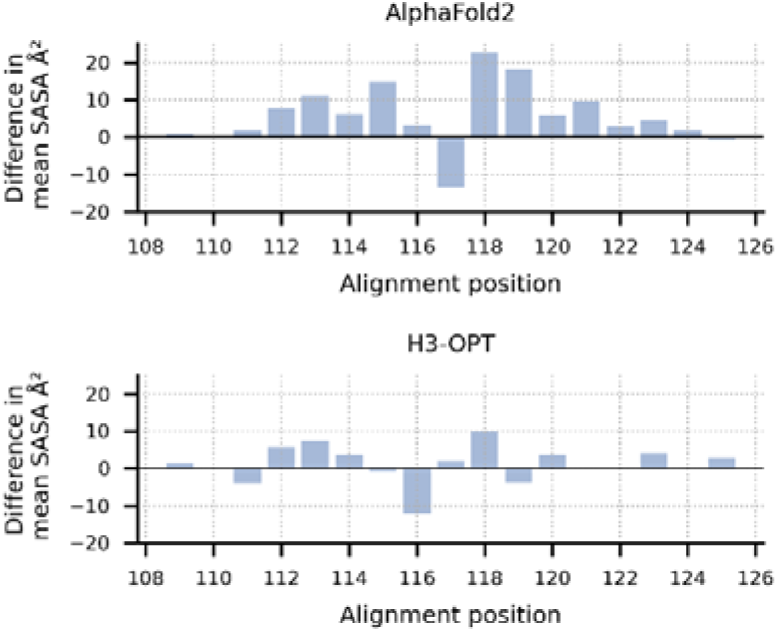
Solvent-accessible surface area analysis of predicted H3 loops. The values represent the difference in SASA between H3 structures predicted by AF2 or H3-OPT and experimentally determined structures. Positive values indicate that the predicted structures have more exposed surface area compared to the native structures; negative values indicate less exposed surface area.

**Extended Data Fig. 2.**
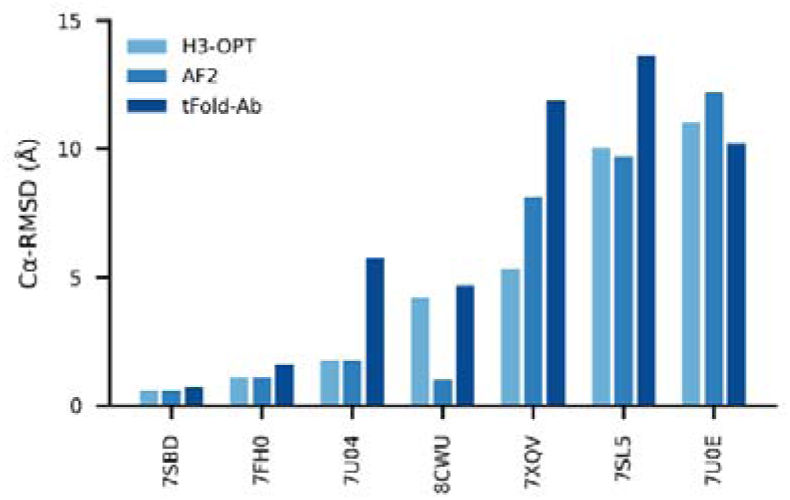
Comparison of accuracy between AF2, H3-OPT, and tFold-Ab methods using the CAMEO 2022 benchmark dataset. The x-axis represents different targets; y-axis represents Cα-RMSD values.

