## Supplemental information for "H3-OPT: Accurate prediction of CDR-H3 loop structures of antibodies with deep learning"

[1.3 AlphaFold2 accurately predicted the V](#_Toc141039543)_[H](#_Toc141039543)_[/V](#_Toc141039543)_[L](#_Toc141039543)_ [orientations and CDR-H3 loops of antibodies.	4](#_Toc141039543)

### 1. Supplementary analysis of AlphaFold2 on antibody modeling

#### 1.1 High-quality antibody crystal structures in benchmarks.

To evaluate the performance of AlphaFold2^1^ (AF2) against currently available methods, we conducted two datasets (DB1 and DB2) with high-resolution (< 2.5 Å) X-ray crystal structures from SAbDab^2^. The CDR length distributions were similar in each dataset (Fig. S1b). Additionally, we plotted sequence logos and identified the CDR loops of all heavy chain sequences (Fig. S1c-d). These sequence logo plots revealed higher degree of sequence variability of CDR loops, particularly in the CDR-H3, with smaller residue letters than framework regions. Overall, high-quality and diverse antibody datasets allowed us to assess the quality of predicted models and evaluate the strengths and weaknesses of all alternative methods.

#### 1.2 Accuracy of AF2 for the overall antibody structure predictions was remarkable.

To assess the similarity between the predicted models generated by each method and the experimentally determined structure, we first computed the template modeling scores (TM-scores)^3^ for all predicted models of each target. The TM-scores of AF2 predictions in different datasets were as follows: DB1, 0.93 ± 0.04; DB2, 0.94 ± 0.03 (Fig. S1e). Although the average TM-score of AF2 was slightly lower than those of DeepAb^4^ in DB1, it outperformed DeepAb by a large margin in DB2 (0.94 vs 0.87, on average). Then, we used global distance test (GDT) scores^5^ to assess the similarity of predicted substructures at different structural cutoffs. The average GDT-TS scores of the models generated by AF2, AbodyBuilder^6^, and DeepAb in DB1 were 0.90, 0.88, and 0.91, respectively. In DB2, the GDT-TS scores of AF2, RoseTTAFold^7^, and NanoNet^8^ all exceeded 0.90. Furthermore, we found that the models with high GDT-TS scores also exhibited higher GDT-HA scores compared to the models with low GDT-TS scores. We finally calculated the average Z-scores of GDT-TS, GDT-HA, and TM-score for all datasets to provide comprehensive scores for each method. AF2 outperformed other methods on both DB1 and DB2, with an average Z-score of 0.87 and 1.08, respectively.

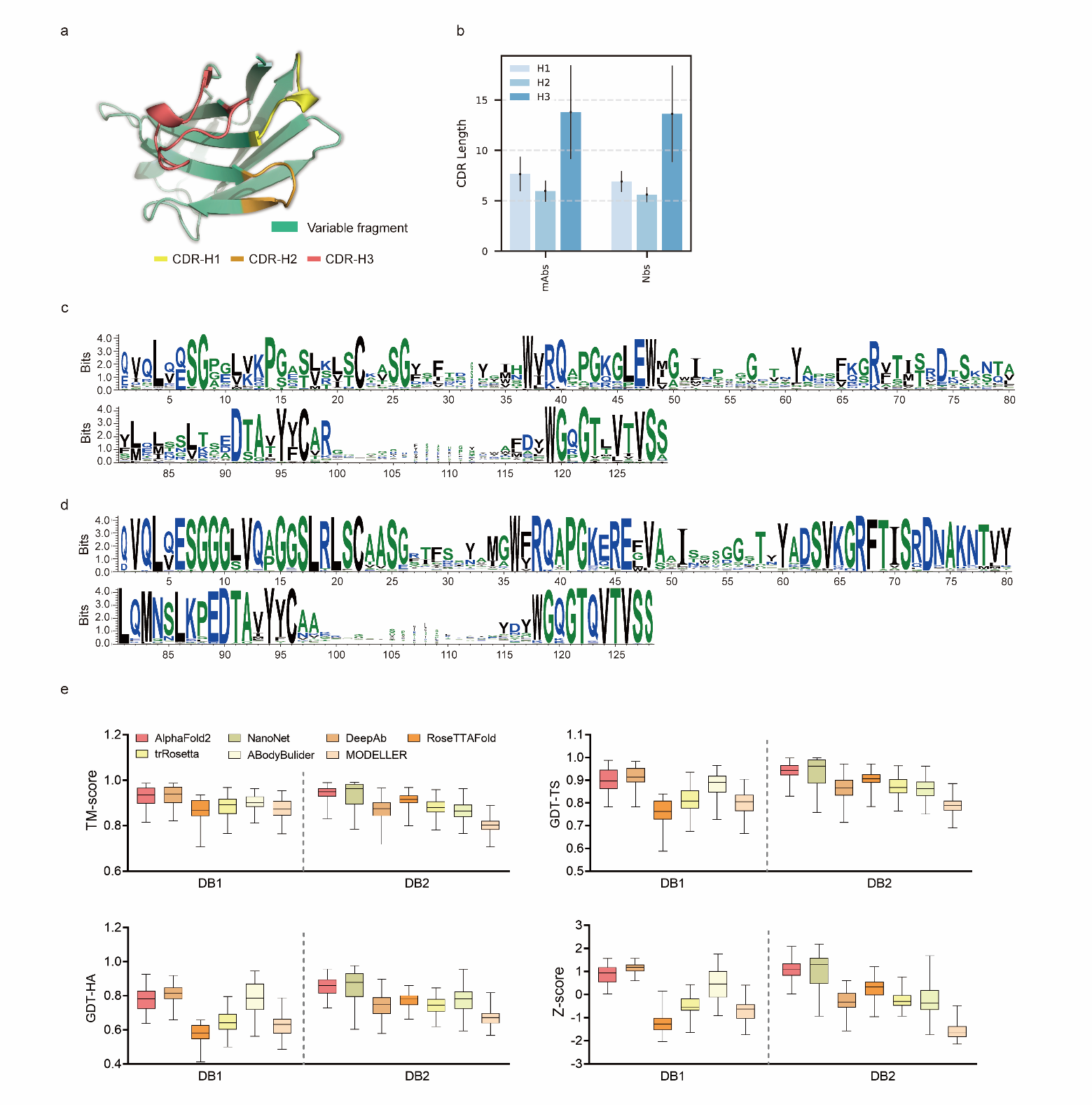

**Figure S1| Accuracy of AF2 on antibody modeling. a**, Schematic for CDR heavy chain loops. **b,** The CDR lengths of mAbs (n = 47) and Nbs (n = 78). The error bars represent the standard deviation of the data. **c,** Sequence logo plots of V_H_ fragments in DB1. **d,** Sequence logo plots of V_H_ fragments in DB2. The different colors of codes represent the hydrophobicity of amino acids. **e,** The performance of AF2 on different datasets using various evaluation metrics. In the box plots, the lines at the center of the boxes represent the medians, and the top and bottom lines of the boxes represent the upper and lower quartiles.

#### 1.3 AlphaFold2 accurately predicted the V_H_/V_L_ orientations and CDR-H3 loops of antibodies.

Given the vital roles of CDR-H3 loops in antigen recognition, we assessed the local accuracy of CDR-H3 loops among all participants. We first superimposed the backbone atoms of the entire Fv regions to reference structures and calculated the heavy atom RMSDs of CDR-H3 loops (RMSD_HA_). The average RMSD_HA_ values of AF2, DeepAb and AbodyBuilder in DB1 were 3.92, 3.64 and 3.69 Å, respectively, which were lower than those of other methods (Fig. S2a). Additionally, these RMSD_HA_ values decreased slightly after superimposing the V_H_ backbone heavy atoms, indicating that these methods could predict V_H_/V_L_ orientations more accurately than other alternative methods (Fig. S2b). Similar to TM-score, DeepAb performed better than AF2 in DB1, but its accuracy decreased substantially in DB2 (Fig. S2c). AF2, NanoNet and AbodyBuilder outperformed other methods in DB2, with average RMSD_HA_ values of 3.79, 3.44 and 4.37 Å, respectively.

To better understand the prediction results, we next conducted side-to-side comparisons for above-mentioned methods. The results showed that there were no significant differences in either the backbone or CDR-H3 RMSDs after superimposing the V_H_ backbone heavy atoms between DeepAb and AF2 (Fig. S3a). Additionally, AF2 accurately produced the Nb structures with lower backbone RMSDs (all below 4Å), but provided comparative accuracy in CDR-H3 RMSDs with NanoNet (Fig. S3b). We found that the main reason for this result was that NanoNet predicted wrong main-chain structures at the C-terminus. AbodyBuilder utilized homology modeling-based algorithm for structure prediction, providing similar or better backbone quality than AF2 (Fig. S3c-d). However, it did not significantly improve the accuracy of the CDR3 loops because AbodyBuilder highly relied on the quality of templates (especially when modeling DB2). Taken together, these results indicated that AF2 was a highly effective tool for predicting antibody structures both in mAbs and Nbs, and it produced CDR-H3 loops that were comparable to those generated by AI-based antibody specific methods.

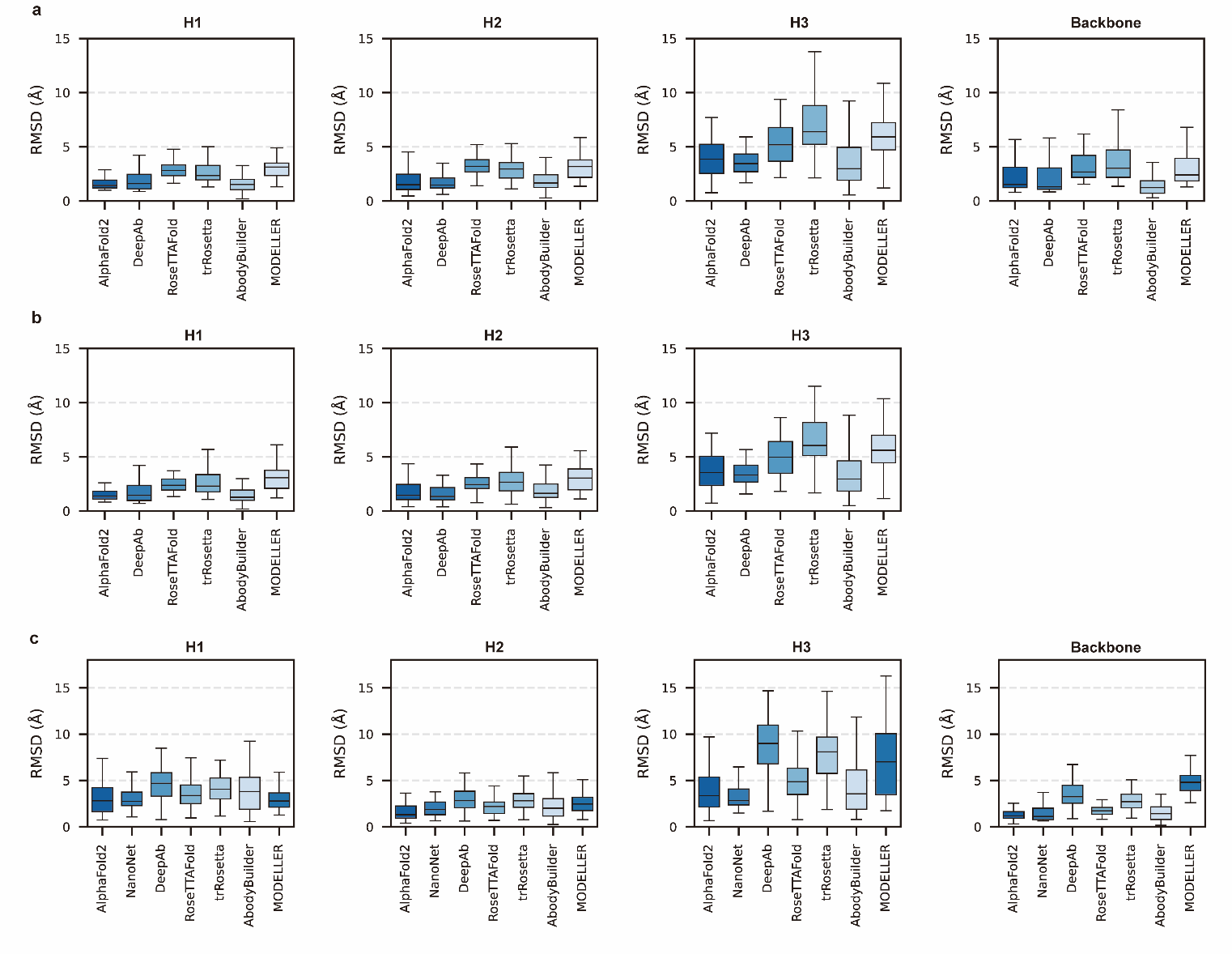

**Figure S2| Accuracy of AF2 on different antibody regions. a,** The performance of AlphaFold2 in DB1 relative to other methods after superimposing Fv backbones. **b,** The performance of H3-OPT in DB1 relative to other methods after superimposing V_H_ backbones. **c,** The performance of H3-OPT in DB2 relative to other methods.

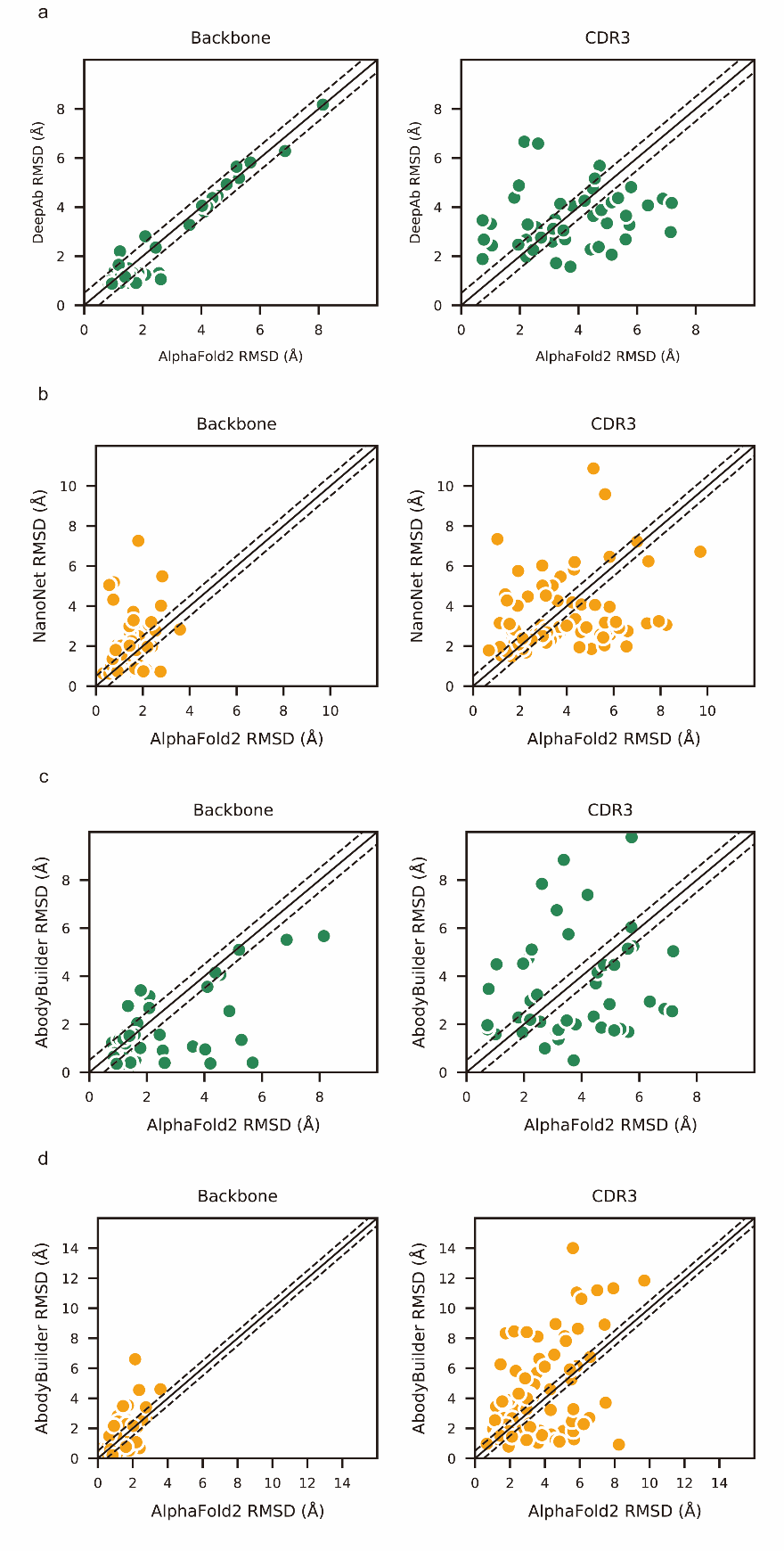

**Figure S3| Local accuracy of AlphaFold2 prediction. a,** Side-by-side comparison of Backbone and CDR3 heavy atom RMSDs for DeepAb and AlphaFold2 in DB1. **b,** Side-by-side comparison of Backbone and CDR3 heavy atom RMSDs for NanoNet and AlphaFold2 in DB2. **c,** Side-by-side comparison of Backbone and CDR3 heavy atom RMSDs for AbodyBuilder and AlphaFold2 in DB1. **d,** Side-by-side comparison of Backbone and CDR3 heavy atom RMSDs for AbodyBuilder and AlphaFold2 in DB2. The slashes represent a cutoff value of 0.5 Å.

### 2. Supplementary analysis of computational approaches on H3 optimization

#### 2.1 Quantum mechanics-based optimization failed to optimize CDR-H3 loops.

Due to the lack of consideration of electronic effects in AF2, we hypothesized that the accuracy of CDR-H3 could be further improved by quantum mechanics (QM)-based methods. We applied two QM-based approaches to optimize AF2 models: the first was an energy-based re-ranking method, and the second was loop optimization with QM. For the first approach, we introduced a new dataset containing 14 antibody sequences with the same CDR-H3 and then run QM optimization for all AF2 models. These models were re-ranked according to the QM energies. As shown in Table 1, our results demonstrated that this method outperformed the default ranking criteria for 8 out of 14 targets, achieving an average RMSD improvement of 0.69 Å. However, this re-ranking method did not yield more accurate results when applied to a larger database (DB3) (Table 2). We also made additional attempts to improve the re-ranking process by removing constraints on terminal atoms, altering re-ranking criteria, and considering solvent effects. However, these methods did not improve the accuracy of AF2 (ΔRMSDs < 0). Moreover, the QM-based optimization methods also failed to improve the accuracy of H3 loops (Table 3). We finally generated structures based on the conformational proportions using Boltzmann distribution of the QM energies, but the accuracy of these structures did not match that of Ranked_0, with all ΔRMSD below 0. In conclusion, although QM-based methods may not improve the accuracy of CDR-H3 loops in general cases, it still provided opportunities for loop modeling as the development of more accurate physics-based methods.

Table 1. The RMSD results of PM6D3 level re-ranking method on 14 same CDR-H3 antibodies

| **PDBID** | | **Ranked 0 RMSD** | **Lowest energy RMSD** | | **Lowest RMSD** | **Δ_RMSD_*** |
| --- | --- | --- | --- | --- | --- | --- |
| **4kmt** | 1.06 | | | 1.14 | 1.05 | -0.08 |
| **5i19** | 2.16 | | | 1.91 | 1.77 | 0.25 |
| **5i1l** | 3.80 | | | 3.20 | 3.19 | 0.60 |
| **5i17** | 2.86 | | | 3.71 | 2.86 | -0.85 |
| **5i1d** | 2.10 | | | 2.10 | 2.02 | 0.00 |
| **5i1c** | 2.43 | | | 1.66 | 1.45 | 0.77 |
| **5i1a** | 2.37 | | | 0.85 | 0.59 | 1.52 |
| **5i1i** | 3.72 | | | 3.51 | 3.51 | 0.21 |
| **5i15** | 2.16 | | | 1.94 | 1.35 | 0.22 |
| **5i16** | 3.19 | | | 1.70 | 1.39 | 1.49 |
| **5i18** | 2.88 | | | 2.88 | 2.88 | 0.00 |
| **5i1e** | 1.62 | | | 1.13 | 0.92 | 0.49 |
| **5i1g** | 2.08 | | | 2.08 | 2.00 | 0.00 |
| **5i1h** | 1.58 | | | 1.84 | 1.32 | -0.26 |

*Δ_RMSD_ was calculated by subtracting the RMSD of predicted model from the RMSD of Ranked_0 model.

Table 2. Accuracy of QM-based re-ranking methods

| **Method** | **Freeze terminal Cα** | **CDR*** | **Phase** | **Ranked 0 RMSD** | **Lowest energy RMSD** | **Lowest RMSD** | **Δ_RMSD_** |
| --- | --- | --- | --- | --- | --- | --- | --- |
| PM6D3 | Y | H3 | Gas | 2.64 | 2.76 | 2.16 | -0.12 |
| PM6D3 | N | H3 | Gas | 2.53 | 2.67 | 2.03 | -0.14 |
| PM6D3 | N | H1, H2, H3 | Gas | 2.50 | 2.64 | 2.00 | -0.14 |
| B3LYP | N | H3 | Gas | 2.66 | 2.87 | 2.30 | -0.21 |
| B3LYP | N | H3 | Water | 2.66 | 2.68 | 2.30 | -0.02 |

*CDR means the energy of which loop is used to re-rank AF2 models.

Table 3. Accuracy of QM-based optimization methods

| **Method** | **Freeze terminal Cα** | **Structure generation method** | **Phase** | **Ranked 0 RMSD** | **Lowest energy RMSD/Opted RMSD** | **Lowest RMSD** | **Δ_RMSD_** |
| --- | --- | --- | --- | --- | --- | --- | --- |
| PM6D3 | Y | / | Gas | 1.69 | 1.74/1.87 | 1.37 | -0.05 |
| B3LYP | N | / | Gas | 1.63 | 1.65/2.55 | 1.38 | -0.02 |
| B3LYP | N | / | Water | 1.63 | 1.58/2.25 | 1.38 | 0.05 |
| B3LYP | N | Boltzmann | Gas | 1.56 | 2.05 | 1.28 | -0.49 |
| B3LYP | N | Boltzmann | Water | 1.56 | 1.81 | 1.28 | -0.25 |
| B3LYP | N | Boltzmann, minimized | Gas | 1.56 | 1.96 | 1.28 | -0.40 |
| B3LYP | N | Boltzmann, minimized | Water | 1.56 | 1.84 | 1.28 | -0.28 |

#### 2.2 Molecular dynamics (MD) simulations could not provide accurate CDR-H3 loop conformations.

MD simulations were extensively used to explore stable conformations of proteins in a water environment^9^ and were also introduced to loop modeling^10^. We used MD simulations to search for representative CDR-H3 loop conformations. The simulation systems consist of antibody structures, water and ions. We selected the top 10 challenging-to-predict targets from the DB1 and DB2 to investigate the dynamic conformations of CDR-H3 loops. The first target was 7N0R, a single-domain antibody that binds the SARS-CoV-2 Nucleocapsid protein with a 12-residue CDR-H3 loop^11^. We found that MD successfully generated conformations with lower CDR-H3 Cα-RMSDs (average value is 5.62 Å, achieving an improvement of 5.30 Å over the AF2 prediction) by correcting the orientation of the CDR-H3 loop (Fig. S4a). Additionally, the conformations of 7N0R indicated that the CDR-H3 loop was more flexible than framework regions and other CDR loops, demonstrated by root mean square fluctuation (RMSF) of every residue during simulation (Fig. S4b). Despite these promising results for 7N0R, the high CDR-H3 Cα-RMSDs of all 10 targets (> 5 Å) and poor performance in other cases (ΔCα-RMSD < 1.5 Å) (Table 4) both suggested that the conformation search method based on MD simulations failed to generate CDR-H3 loops that closely matched native structures. However, the accuracy of these antibodies with relative long CDR-H3 loops could be improved by H3-OPT through incorporating latent structural information of loop folding from ESM2 or enhanced sampling techniques^12^.

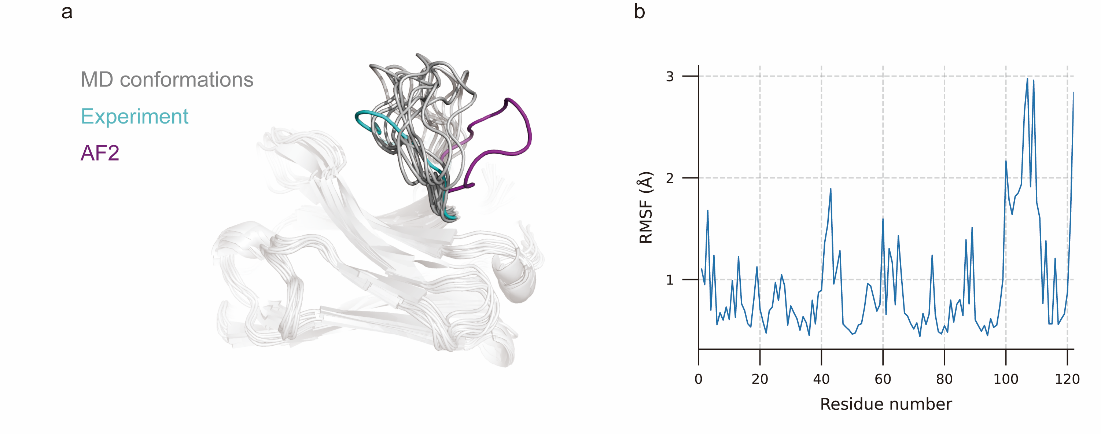

**Figure S4| MD generated conformations for benchmark target 7N0R. a,** Comparison of CDR-H3 loops of MD (gray), AF2 (pink) and experimentally determined structure (cyan). **b,** RMSF of antibody residues during simulation. CDR-H3 loop is located in residue number ranging from 98 to 109.

Table 4. The accuracy of MD-based CDR-H3 loop optimization in the 10 worst cases of AF2

| PDBID | Cα-RMSD_Ranked_0_ | Cα-RMSD_MD_opt_ | ΔCα-RMSD |
| --- | --- | --- | --- |
| 7n0r | 10.92 | 5.62±0.97 | 5.30 |
| 3juy | 6.37 | 5.71±0.23 | 0.66 |
| 5y80 | 6.61 | 7.59±0.47 | -0.98 |
| 7a4t | 6.19 | 7.48±0.29 | -1.29 |
| 4nzr | 6.57 | 7.73±0.26 | -1.16 |
| 6xzu | 7.45 | 6.34±0.94 | 1.11 |
| 6x05 | 6.32 | 7.48±0.63 | -1.16 |
| 3c08 | 6.68 | 7.01±0.11 | -0.33 |
| 4z9k | 9.04 | 8.01±0.37 | 1.03 |
| 6oca | 7.61 | 8.01±0.34 | -0.40 |

### 3. Supplementary methods

**Datasets**

High-quality structures are necessary for the accurate assessment of antibody modeling methods. All antibody information was obtained from the SAbDab website. We first downloaded antibody structures with X-ray diffraction resolution of 2.5 Å or less from the RCSB website (<https://www.rcsb.org/>) and deleted redundant sequences with 95% or more identity. Then, we clustered the remaining sequences into 93 clusters using UCLUST^13^ by 65% sequence similarity cutoff and selected centroid sequences to ensure the representativeness and sufficient quantity of the final datasets. We additionally omitted the PDBIDs 5DHV, 3SE9, 4YDJ, 5VF6, 6CT7, 7FAB, 6NMT, 5ILT, and 5UXQ which were unable to predict using AbodyBuilder. The Fv sequence of each target was identified from initial sequences using the Chothia^14^ definitions. For the Nb dataset, we collected Nb sequences from a non-redundant data set^15^ and deleted all structures with low resolution (≥ 2.5 Å) to construct a high-resolution nanobody structure database. Seven native structures (PDBIDs 6NEX, 6FFJ, 4BUH, 6BHZ, 6UUM, 5O03, and 5C1M) were further excluded from the final dataset owing to the missing residues in CDR-H3 loops. In total, 125 targets were included in the final datasets: 47 mAbs (DB1) and 78 Nbs (DB2). After the model assessment, additional 98 antibodies and 42 nanobodies, which were collected from the test sets of DeepAb and NanoNet, were combined to verify the improvement of the CDR-H3 optimization methods (DB3).

**Sequence logos generation**

Sequences of DB1, DB2 V_H_ domains were determined by Chothia definitions and aligned using ANARCI^16^ with the Aho scheme^17^. Then, we removed the alignment positions with 85% gaps to obtain a good visualization of alignment statistics. Finally, the alignment outputs of different datasets were submitted to the WebLogo3 server^18^ to obtain sequence logos of each database. These sequence logos quantitatively showed the conservation of antibody sequences. The height of each letter within the logo corresponds to its base frequency at that particular position. The letters are arranged in descending order of size, with the tallest (or most frequent) letter positioned at the top.

**Benchmarking alternative methods** To evaluate the prediction accuracy of all alternative methods, we benchmarked five AI-based methods (AF2, RoseTTAFold, trRosetta^19^, NanoNet, and DeepAb) and two TBM methods (MODELLER^20^ and ABodyBuilder). Identical templates were excluded from TBM templates database and AF2 PDB database. To model full-length V_H_ and V_L_ antibody, fifty glycines were added as a linker between two chains. The output multiple sequence alignment files of AF2 were then used to predict MODELLER models. ABodyBuilder models were generated using the webserver and modelled by Sphinx. For all methods, recommended parameters were used for all antibody predictions. NanoNet was removed from the DB1 as it was only applied for Nb modeling. We only compared and summarized the target structures which were successfully produced by all participants.

**Model assessment**

The residue indexes of different models were aligned and renumbered using MAFFT ^21^ to obtain the same sequence length. We assessed the accuracy of AF2 through varying metrics, including TM-score, GDT-TS score and GDT-HA score. The structural cutoffs for GDT-TS score were 1, 2, 4 and 8 Å, respectively while that GDT-HA score were 0.5, 1, 2, and 4 Å, respectively. We calculated the average Z-scores of these methods to assess the overall similarities. The average Z-score of each target was given by:

$$Z_{ave}=\frac{1}{3}Z_{GDT-TS}+ \frac{1}{3}Z_{GDT-HA}+\frac{1}{3}Z_{TM-score}$$

To better evaluate the accuracy of CDR loops, RMSDs were computed after superimposing the Fv or V_H_ backbone heavy atoms to their reference structures. All RMSDs were computed using Schrödinger v. 2017-2 (Schrödinger, New York, NY, USA).

**Molecular dynamic simulations**

We analyzed the binding affinities of antibody-antigen complexes predicted by AF2 and H3-OPT. Their relative binding affinities were calculated through molecular dynamics simulations. Initially, the missing side-chains and loops in the antibody structures were filled using the Protein Preparation Wizard module within Schrödinger software. Subsequently, the tLEaP module of AMBER^22^ was employed to construct the simulation system with the ff19SB force field^23^ and OPC solvent model^24^. Additionally, the simulation system was solvated with a 0.15 M NaCl solution. Energy minimization was performed through a 5000-step steepest descent algorithm, followed by a 5000-step conjugate gradient algorithm. Afterwards, a 25000-step NVT simulation with a time step of 1 fs was performed to gradually heat the system from 0 K to 100 K. A 250000-step NPT simulation with a time step of 2 fs was carried out to further heat the system from 100 K to 298 K. In the production run, 100-ns MD simulations were performed with a time step of 2 fs. The distance cutoff for nonbonded interactions was set to 10 Å, and the Berendsen algorithm was utilized to maintain isotropic pressure coupling at 1 bar. The Langevin algorithm was employed to maintain the simulation temperature at 298 K.

**Refinement of the CDR-H3 loop of AF2 through quantum-mechanics (QM) methods**

We extracted all H3 loop atoms from different AF2 models re-rank these models based on QM energies. The nitrogen atoms and carbonyl atoms were initially removed from N-terminals and C-terminals, respectively. Next, we added hydrogens to these loops and produced the input files for different QM optimization methods using Schrödinger v. 2017-2. The geometry optimizations were carried out with the Gaussian16^25^ software. We first optimized the CDR-H3 loop by using PM6-D3^26,27^ and B3LYP/6-31G*^28,29^. The optimization method was divided into three steps: (1) optimizing hydrogens; (2) freezing the terminal C-alpha atoms and optimizing other atoms; (3) frequency calculations to obtain Gibbs free energies. For an alternative CDR-H3 loop optimization method, we calculated the single point energies with thermal corrections. The solvation effect can be estimated using the polarizable continuum model (with a dielectric constant ε=78.36). After optimization, the proportion of each conformation was calculated based on energies of five AF2 models according to Boltzmann distribution, which was used to generate a Boltzmann-averaged structure. The structures which failed in the optimization process were excluded from the final comparison.

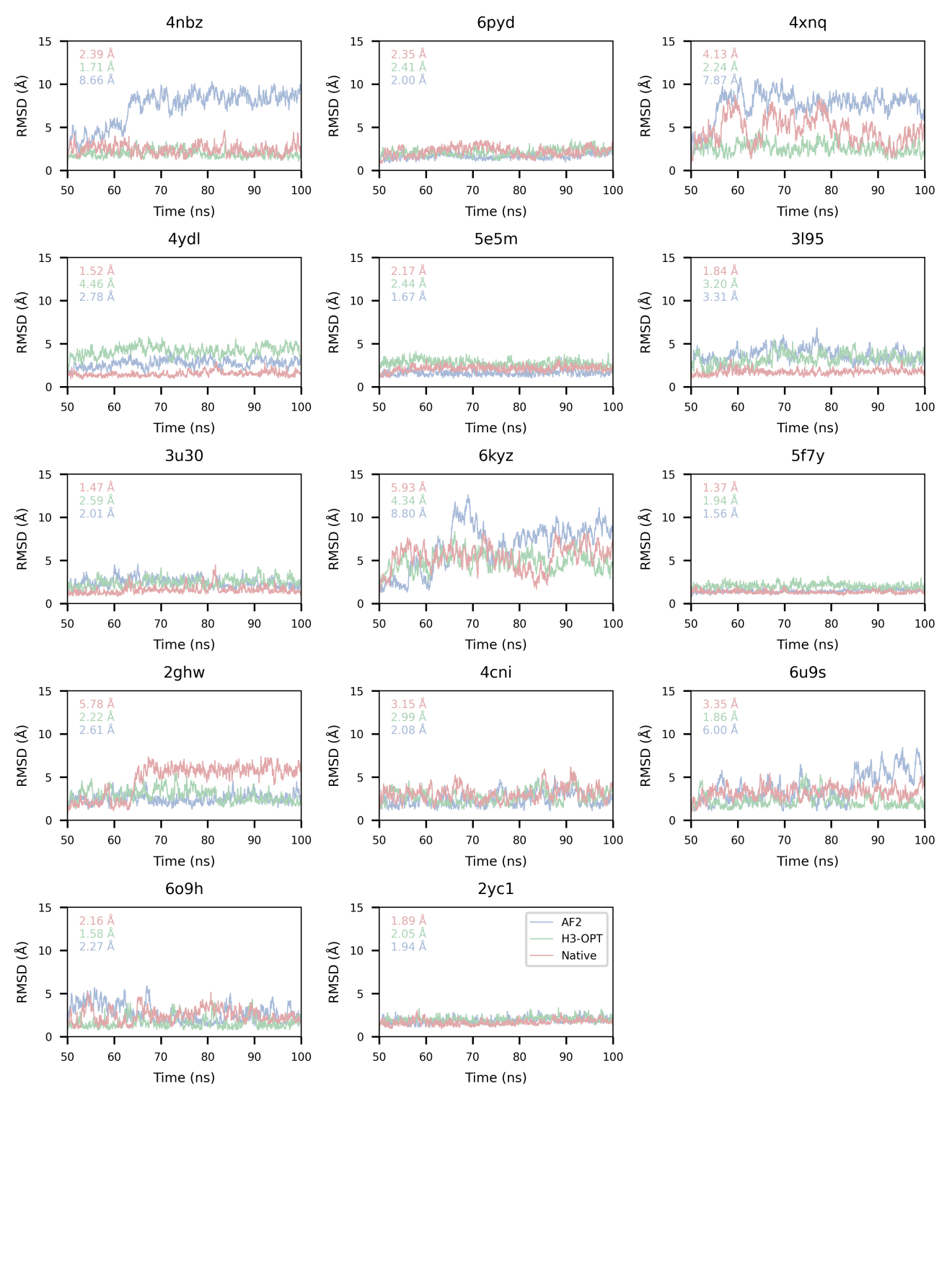

**Figure S5| RMSDs_backbone_ of 14 complexes during MD simulations.** The average RMSD values of the backbone for the last 10 nanoseconds are presented in the top-left corner.
